# Reticular Thalamic Hyperexcitability Drives Autism Spectrum Disorder Behaviors in the Cntnap2 Model of Autism

**DOI:** 10.1101/2025.03.21.644680

**Authors:** Sung-Soo Jang, Fuga Takahashi, John R Huguenard

## Abstract

Autism spectrum disorders (ASDs) are a group of neurodevelopmental disorders characterized by social communication deficits, repetitive behaviors, and comorbidities such as sensory abnormalities, sleep disturbances, and seizures. Dysregulation of thalamocortical circuits has been implicated in these comorbid features, yet their precise roles in ASD pathophysiology remain elusive. This study focuses on the reticular thalamic nucleus (RT), a key regulator of thalamocortical interactions, to elucidate its contribution to ASD-related behavioral deficits using a Cntnap2 knockout (KO) mouse model. Our behavioral and EEG analyses comparing Cntnap2^+/+^ and Cntnap2^-/-^ mice demonstrated that Cntnap2 knockout heightened seizure susceptibility, elevated locomotor activity, and produced hallmark ASD phenotypes, including social deficits, and repetitive behaviors. Electrophysiological recordings from thalamic brain slices revealed increased spontaneous and evoked network oscillations with increased RT excitability due to enhanced T-type calcium currents and burst firing. We observed behavior related heightened RT population activity in vivo with fiber photometry. Notably, suppressing RT activity via Z944, a T-type calcium channel blocker, and via C21 and the inhibitory DREADD hM4Di, improved ASD-related behavioral deficits. These findings identify RT hyperexcitability as a mechanistic driver of ASD behaviors and underscore RT as a potential therapeutic target for modulating thalamocortical circuit dysfunction in ASD.

**Teaser:** RT hyperexcitability drives ASD behaviors in Cntnap2-/- mice, highlighting RT as a therapeutic target for circuit dysfunction.

## Introduction

Autism spectrum disorders (ASDs) are common neurodevelopmental conditions characterized by social impairments, repetitive behaviors, and comorbidities such as intellectual disability, hyperactivity, anxiety, and epilepsy. Extensive research has sought to unravel its pathophysiology by investigating cellular and circuit-level mechanisms across multiple brain regions, including the hippocampus, prefrontal cortex, and medial septum, using various ASD-associated risk genes and animal models (*1–9*). Notably, individuals with ASD often exhibit sensory processing abnormalities (*10*, *11*), sleep disturbances (*12*, *13*), and seizures (*14*, *15*), implicating a potential role of the thalamocortical circuit, a key system governing these functions. Structural alterations in the thalamus and atypical thalamocortical connectivity have been reported in both juvenile and adult individuals with ASD (*16–18*). Recent findings further establish a link between electrophysiological abnormalities within the somatosensory thalamic nucleus and the sensory hypersensitivity and sleep fragmentation observed in children with ASD (*19*).

The reticular thalamic nucleus (RT) is a shell-like GABAergic nucleus with inhibitory projections targeted exclusively to the thalamus. There is evidence for heterogeneity of RT neurons (*20–22*), with the majority expressing parvalbumin (PV). RT acts as the “gatekeeper” for sensory processing, fear, attention, and seizure regulation through modulating thalamocortical activity (*23–29*). RT neurons exhibit two distinct firing modes (*20*): bursting and tonic. Bursting firing, driven by T-type Ca^2+^ channels (*30*), generates high frequency action potential trains leading to strong inhibitory responses (*31*, *32*) and subsequent rebound burst firing in downstream thalamocortical (TC) neurons. At the network level this then can trigger RT re-excitation, such that reciprocal interaction between RT and TC neurons forms the basis for an intrinsic thalamic oscillator. Reinforcing this thalamic oscillator are reciprocal excitatory connections with cerebral cortex. RT is the main source of intrathalamic inhibition, orchestrating distinct neural rhythms, including sleep spindles, slow oscillations, and gamma rhythms (*33–36*). Disruptions in RT function perturb these dynamics and have been implicated in neuropsychological conditions, including attention-deficit/hyperactivity disorder (ADHD) (*27*), schizophrenia (*37*, *38*), and depressive-like behaviors (*39*). While emerging evidence implicates RT dysfunction in ASD pathophysiology, the precise cellular and circuit-level mechanisms remain largely unexplored, warranting further investigation.

Cntnap2, contactin-associated protein 2, is implicated in ASD, and Cntnap2^−/−^ mice display a number of ASD-related behaviors, including hyperactivity, epileptic seizures, abnormal sleep-wake time and architecture, degraded and unstable sensory coding, and disrupted spatial discrimination (*1*, *4*, *9*, *40*, *41*). Network, cellular, and structural features that have been described so far in these mice include, (1) altered hippocampal gamma oscillations and sharp wave ripples (SWR) with reduced numbers of PV+ interneurons and impaired perisomatic inhibition onto CA1 pyramidal cells (*9*, *42*), (2) disturbed oscillatory population activity in the medial prefrontal cortex (mPFC) with reduction in the balance of excitation to inhibition (E/I) in L2/3 Pyramidal neurons and spines and synapses (*8*), (3) transient reduction in dendritic spine density and a greater rate of microglial pre-synaptic engulfment in the Anterior Cingulate cortex (*43*).

In this study, we systematically investigated whether RT neurons exhibit electrophysiological alterations in Cntnap2^−/−^ mice and how these changes impact thalamocortical circuit function and behavior. To assess the causal relationship between RT dysfunction and ASD-related behavioral deficits, we employed pharmacological and chemogenetic approaches to suppress RT activity. Both interventions successfully ameliorated behavioral impairments, albeit through distinct mechanisms. These findings provide the first evidence of RT hyperexcitability as a contributor to ASD-related deficits and identify RT as a potential therapeutic target for modulating thalamocortical circuit dysfunction in ASD.

## Results

### Elevated seizure susceptibility and locomotor activity in Cntnap2^−/−^ mice

Previous studies have reported a comorbidity between autism spectrum disorder (ASD), hyperactivity, and absence-type seizures in both patients and animal models (*40*, *45*). To investigate whether Cntnap2^−/−^ mice exhibit altered seizure susceptibility and locomotor activity, we monitored behavior and cortical activity using electroencephalogram (EEG) recordings before and following intraperitoneal (i.p.) administration of pentylenetetrazole (PTZ), a GABA_A_ receptor antagonist. After a low dose of PTZ (20 mg/kg) that in mice is generally either subthreshold for evoking seizures or only produces mild, non-convulsive seizures (*46–50*), Cntnap2^-/-^ mice exhibited a range of seizure phenotypes, including spike-wave discharges (SWDs) at 4–7 Hz, myoclonic jerks (MJs), generalized tonic-clonic seizures (GTCS), and occasional mortality during the 10-minute monitoring period (Fig. 1A). Seizure progression was common, especially in Cntnap2^−/−^ mice (e.g. Fig. 1A), where initial SWDs were seen, followed by MJs, then GTCSs and death. 6 of 10 Cntnap2^−/−^ mice exhibited such seizure progression, whereas only 4 of 11 Cntnap2^+/+^ mice showed any progression past the initial SWD stage; GTCSs were rare, no animals experienced seizure-related death. Cntnap2^−/−^ mice demonstrated significantly shorter latencies to both MJs (Cntnap2^+/+^ mice: 9/11; Cntnap2^−/−^ mice: 8/10) and GTCS (Cntnap2^+/+^ mice: 5/11; Cntnap2^−/−^ mice: 8/10) compared to Cntnap2^+/+^ controls (Fig. 1, B and C).

**Fig. 1.**
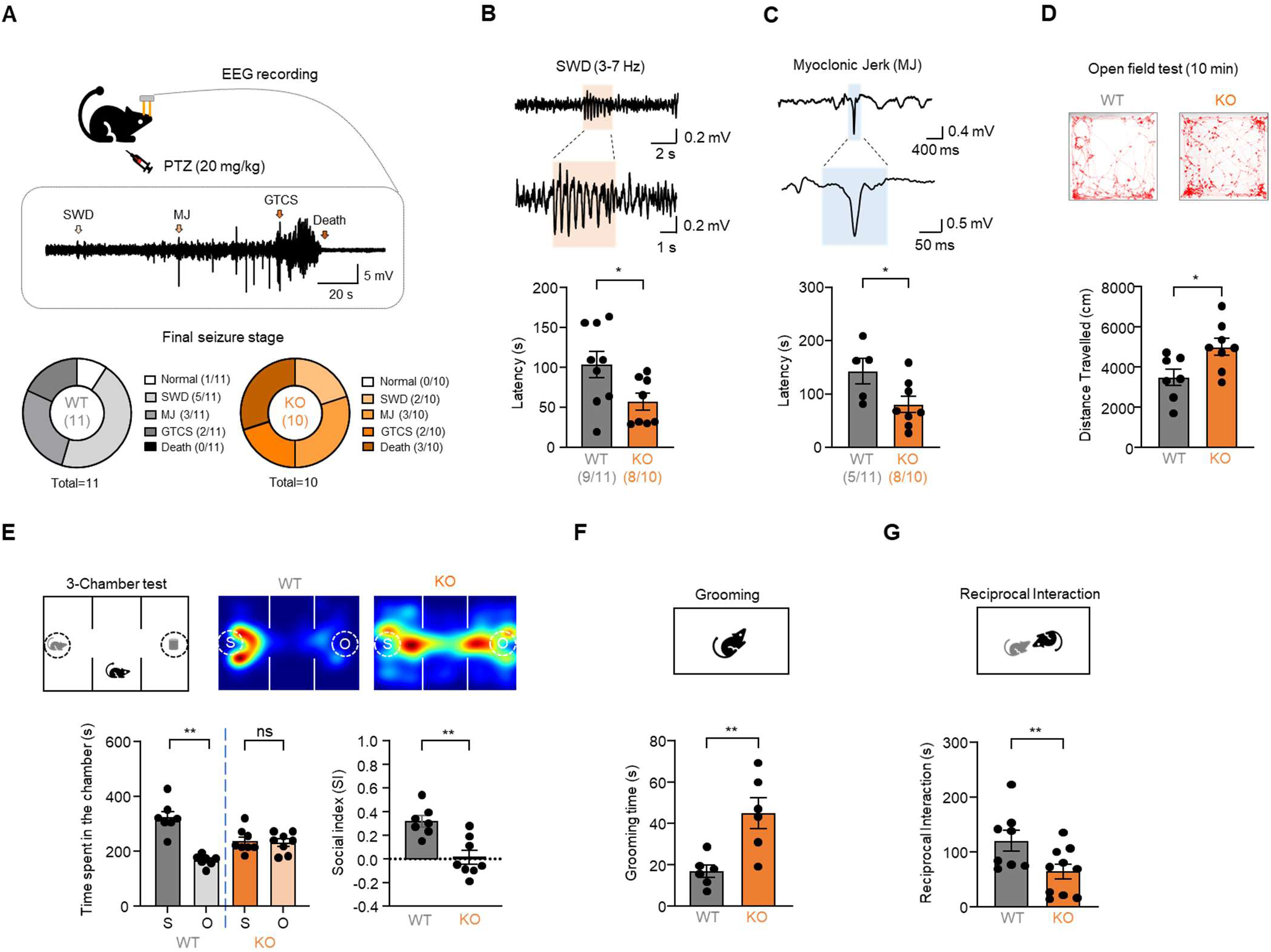
Cntnap2^−/−^ mice display elevated seizure susceptibility, hyper-locomotion, and other ASD-related behaviors. **(A)** Representative examples showing epileptiform spike discharges, including SWD, MJ, GTCS, and death in Cntnap2^−/−^ mice monitored following i.p injection of 20 mg/kg PTZ, as shown by EEG (top). A pie chart illustrating the distribution of mice by their final seizure stages in Cntnap2^+/+^ and Cntnap2^−/−^ mice (bottom). **(B)** Representative traces showing example SWD (top) and summary of latency to first SWD (bottom). (n = 9 mice for Cntnap2^+/+^ and n = 8 mice for Cntnap2^−/−^). **(C)** Representative trace showing MJ (top) and summary of latency to first MJ (bottom). (n = 5 mice for Cntnap2^+/+^ and n = 8 mice for Cntnap2^−/−^). **(D)** Representative trace showing position tracking in an open arena (top) and summary of distance traveled for 10 min (bottom). (n = 7 mice for Cntnap2^+/+^ and n = 8 mice for Cntnap2^−/−^). **(E)** Schematic of 3-chamber test consisting of a 10-min social preference test phase involving social stimulus and non-social stimulus (top left) and heatmaps of the topographical time distribution (top right). Summary of time spent in the social (S) or object (O) chamber (bottom left) and calculated social index (bottom right). (n = 7 mice for Cntnap2^+/+^ and n = 8 mice for Cntnap2^−/−^). **(F)** Schematic of grooming assay, hand scored, blinded (top) and summary of grooming time (bottom) for 10 min. (n = 6 mice for Cntnap2^+/+^ and n =6 mice for Cntnap2^−/−^). **(G)** Schematic for reciprocal interaction (top) and summary of reciprocal interaction time (bottom) for 10 min. (n = 8 mice for Cntnap2^+/+^ and n =10 mice for Cntnap2^−/−^). The statistical tests involved an unpaired two-tailed t-test (B-G). Data are represented as mean values ± SEM. *P*-values in figure panels: *p < 0.05, **p < 0.01.

Cntnap2^−/−^ mice displayed a number of altered behaviors under control conditions, including increased locomotor activity, as evidenced by a greater total distance traveled in the open field during the 10-minute observation period (Fig. 1D). Social behavior assays revealed social deficits consistent with ASD phenotypes: Cntnap2^−/−^ mice showed little social preference, for example spending equal amounts of time in chambers containing a social partner (S) or an object (O) (Fig. 1E), and exhibited a significantly reduced social index (Fig. 1E). Furthermore, Cntnap2^−/−^ mice displayed increased grooming (Fig.1F) and reduced reciprocal interaction time (Fig. 1G) over the 10-minute test period, further highlighting behavioral abnormalities associated with ASD. These findings demonstrate that Cntnap2^−/−^ mice exhibit core ASD-related behavioral deficits, heightened seizure susceptibility, and increased hyperactivity, establishing their utility as a robust model for studying the pathophysiology of ASD and its associated comorbidities.

### Thalamic Circuit Hyperexcitability in Cntnap2^−/−^ mice

Given the relationship between thalamo-cortical dysfunction and behavioral deficits, including Dravet syndrome (*51*) and absence seizures (*52*, *53*), we hypothesized that Cntnap2^−/−^ mice might display the impairment of thalamocortical circuit. The thalamocortical (TC) loop involves reciprocal long-range connections between specific thalamic nuclei (e.g., ventrobasal thalamus, VB) and corresponding cortical regions (e.g., primary somatosensory cortex), with each projection also sending collaterals to the GABAergic reticular thalamic nucleus (RT), which together generate thalamocortical rhythms (Fig. 2, A and B). The intrathalamic network that supports these rhythms is maintained in horizontally sectioned brain slices (*52–55*), in which evoked oscillations can be triggered with extracellular stimuli applied to the internal capsule. Very occasionally, spontaneously occurring oscillations can also be observed in wt slices (*52*, *56*). By contrast, in slices from Cntnap2^−/−^ mice, spontaneous oscillations (n ≥ 10 within a 10 min recording) were commonly observed (7 of 10 slices) with large numbers of bursts per oscillation (Fig. 2C). In addition, evoked oscillations were enhanced in Cntnap2^−/−^ mice (Fig. 2D), with increased oscillation duration and bursts/oscillation, while oscillation frequency remained unchanged (Fig. 2E). To complement the results with electrical stimulation, we utilized Cntnap2^−/−^ mice x Ai32+Ntsr1-Cre mice, which express channelrhodopsin-2 (ChR2) in layer 6 corticothalamic (L6 CT) neurons projecting to thalamic nuclei, including VB and RT (Fig. 2F) and successfully evoked oscillations via a short optogenetic stimulation (3 mW/cm², 1ms) (Fig. 2G). Similar to results with electrical i.c. stimulation, optogenetic stimulation in slices from Cntnap2^−/−^ mice showed increased oscillation duration and bursts/oscillation, but no change in oscillation frequency (Fig. 2H). Altogether, we observed an increase in spontaneous and evoked thalamic oscillations in Cntnap2^−/−^ mice, which may contribute to the pathophysiology of ASD-related deficits.

**Fig. 2.**
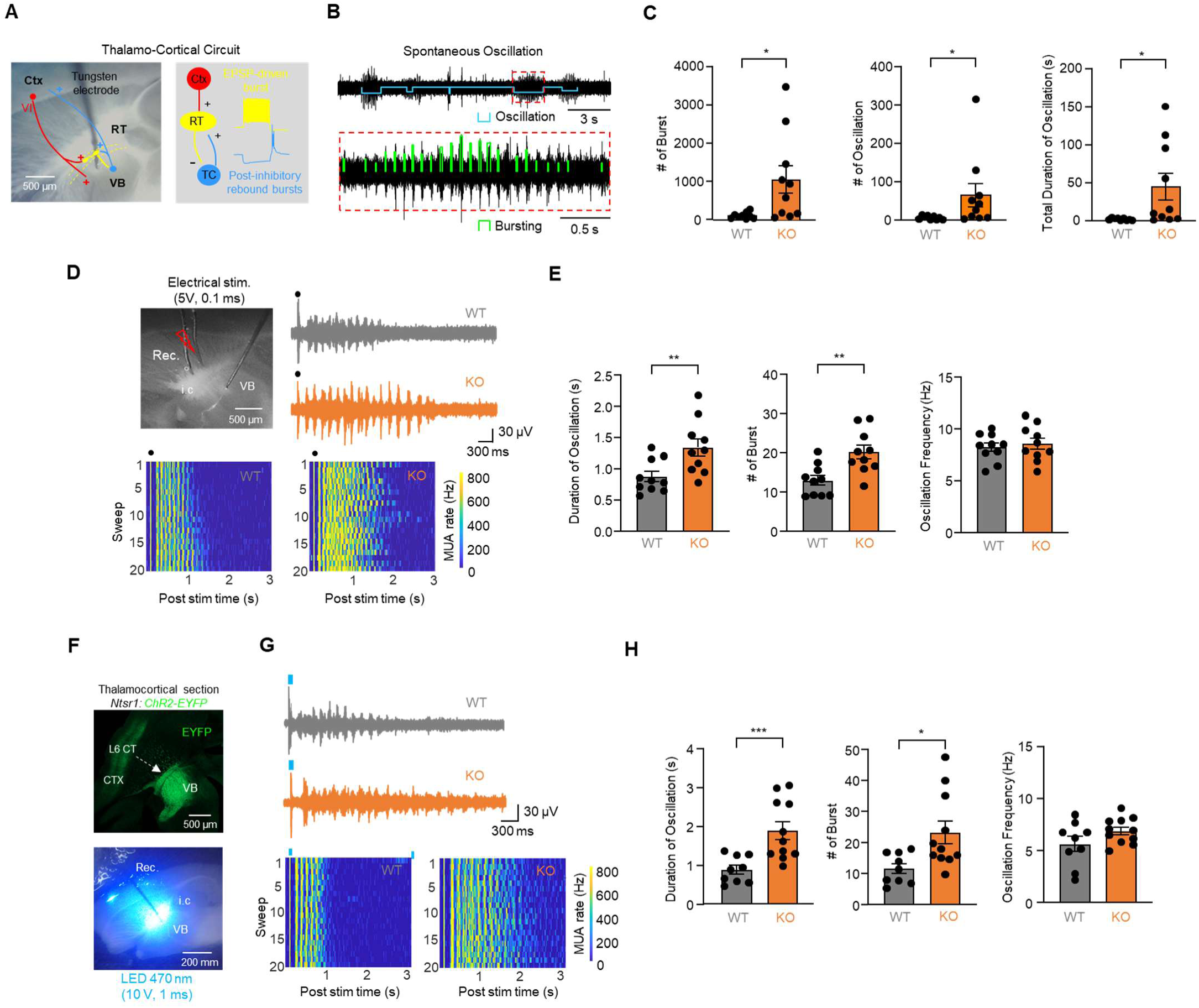
Intra-thalamic circuit is hyperexcitable in Cntnap2^−/−^ mice. **(A)** Schematic depicting on the left the thalamocortical circuit containing corticothalamic cells (red) from layer 6 of somatosensory cortex projecting to VB and RT, the thalamocortical cells (blue) projecting to cortex and sending branches to RT, and RT (yellow) projecting to VB and intra-thalamic connection. Right panel shows membrane responses of RT and TC cells during thalamic oscillations. **(B)** Representative traces showing spontaneous oscillations multiunit recordings of VB cells, with each oscillation (downward blue bracket) composed of many individual bursts of action potentials (upward green brackets). **(C)** Quantification of spontaneous oscillations in terms of bursts per oscillation, number oscillations in 10 minutes, and total time spent oscillating in 10 minutes. (n = 9 slices from 3 mice for Cntnap2^+/+^ and n =10 slices from 3 mice for Cntnap2^−/−^). **(D)** Representative image showing the horizontal thalamic slice preparation with electrical stimulation wires in the internal capsule (i.c.) and a recording electrode (Rec) in VB (top, left). Top right shows representative evoked oscillation in Cntnap2^+/+^ and Cntnap2^−/−^, and bottom heat maps show representative peristimulus time histograms (PSTHs) for detected spikes over 20 sequentially evoked responses. Color intensity codes the number of spikes in each time bin, showing repeated cycles of oscillations lasting for ∼1 s in Cntnap2^+/+^ vs ∼ 2s in Cntnap2^−/−^. **(E)** Quantification of evoked oscillations showing increased bursts and oscillation duration, but not change in frequency. **(F)** Representative image showing the thalamo-cortical section displaying the expression of Ai32-Ntsr1-ChR2-EYFP (Top) and representative image showing the location illuminated by 470 nm LED (bottom). **(G)** Representative traces showing evoked oscillations (top, right), and representative peristimulus time histograms (PSTHs) for example Cntnap2^+/+^ and Cntnap2^−/−^ slices. (H) Quantification of oscillations showing increased bursts and oscillation duration, but not change in frequency. (n = 9 from 4 mice for Cntnap2^+/+^ and n =11 from 4 mice for Cntnap2^−/−^). The statistical tests involved an unpaired two-tailed t-test (B-G). Data are represented as mean values ± SEM. *P*-values in figure panels: *p < 0.05, **p < 0.01.

### Increased burst firing in RT of Cntnap2^−/−^ mice

RT neurons, most of which express parvalbumin (PV) (*57*), provide the primary inhibitory input to thalamocortical (TC) circuits and play a critical role in regulating distinct types of oscillatory activity (*58*). Previous studies have indicated that dysfunction or alterations in PV-expressing neurons render them highly susceptible to factors associated with ASD (*59*, *60*), contributing to deficits such as impaired sensory gating and disruption of the excitation/inhibition balance in the cortex and hippocampus (*61*, *62*). To determine whether electrophysiological properties of RT neurons are altered in Cntnap2^−/−^ mice, we recorded both passive and active properties of RT neurons in brain slices (Fig. 3A). No differences in passive membrane properties including membrane capacitance (pF), resting membrane potential (RMP), and input resistance (MΩ were detected in Cntnap2^−/−^ mice compared to Cntnap2^+/+^ (Fig. S1, A to C). However, the number of action potentials (APs) elicited by depolarizing current injections was significantly increased (Fig. 3B), and this was associated with a reduced rheobase (Fig. 3C). RT responses at rest consisted of an early burst firing phase, complete within 100 ms, and a later tonic firing phase. The number of spikes evoked during the first, bursting phase was increased in Cntnap2^−/−^ mice, but the tonic phase was unaffected (Fig. 3D). No significant changes were observed in individual AP properties, including fast after-hyperpolarization (fAHP), AP threshold, half-width, spike height, max dV/dt, or min dV/dt of the first and last spikes elicited by a 210 pA injection (Fig. S1, D and E). RT cells can fire multiple rebound bursts at the termination of hyperpolarizing steps (*20*, *22*, *51*, *63*). We recorded rebound bursts elicited from a range of holding potentials between –70 to –50 mV. At –70 mV, RT neurons in Cntnap2^−/−^ mice were hyperexcitable, showing rebound bursting, whereas those in Cntnap2^+/+^ mice did not. At –60 mV, most (15 of 18) Cntnap2^−/−^ neurons showed rebound bursts compared to about half (9 of 18) Cntnap2^+/+^ neurons (Fig. 3E). In addition, the number of rebound oscillatory bursts was increased in Cntnap2^−/−^ neurons (Fig. 3F), with no change in the number of spikes in the first rebound burst (Fig. 3G). Low-threshold T-type calcium conductance, which plays a major role in rebound bursting, was significantly increased in Cntnap2^−/−^ neurons. Specifically, the peak T-current density obtained with conditioning potentials in the range of −75 to −105 mV, and test potential of -50 mV, was increased in Cntnap2^−/−^ neurons (Fig. 3H). In contrast to these changes in intrinsic excitability, we observed no differences in synaptic activity; the frequency, amplitude, rise time, or half-width of sEPSCs and sIPSCs of RT neurons were unaffected (Fig. S2, A and B). These findings indicate that Cntnap2^−/−^ neurons exhibit heightened bursting activity, likely driven by increased expression of T-type calcium channels, which likely contributes to heightened thalamocortical circuit oscillations associated with ASD-related deficits.

**Figure 3.**
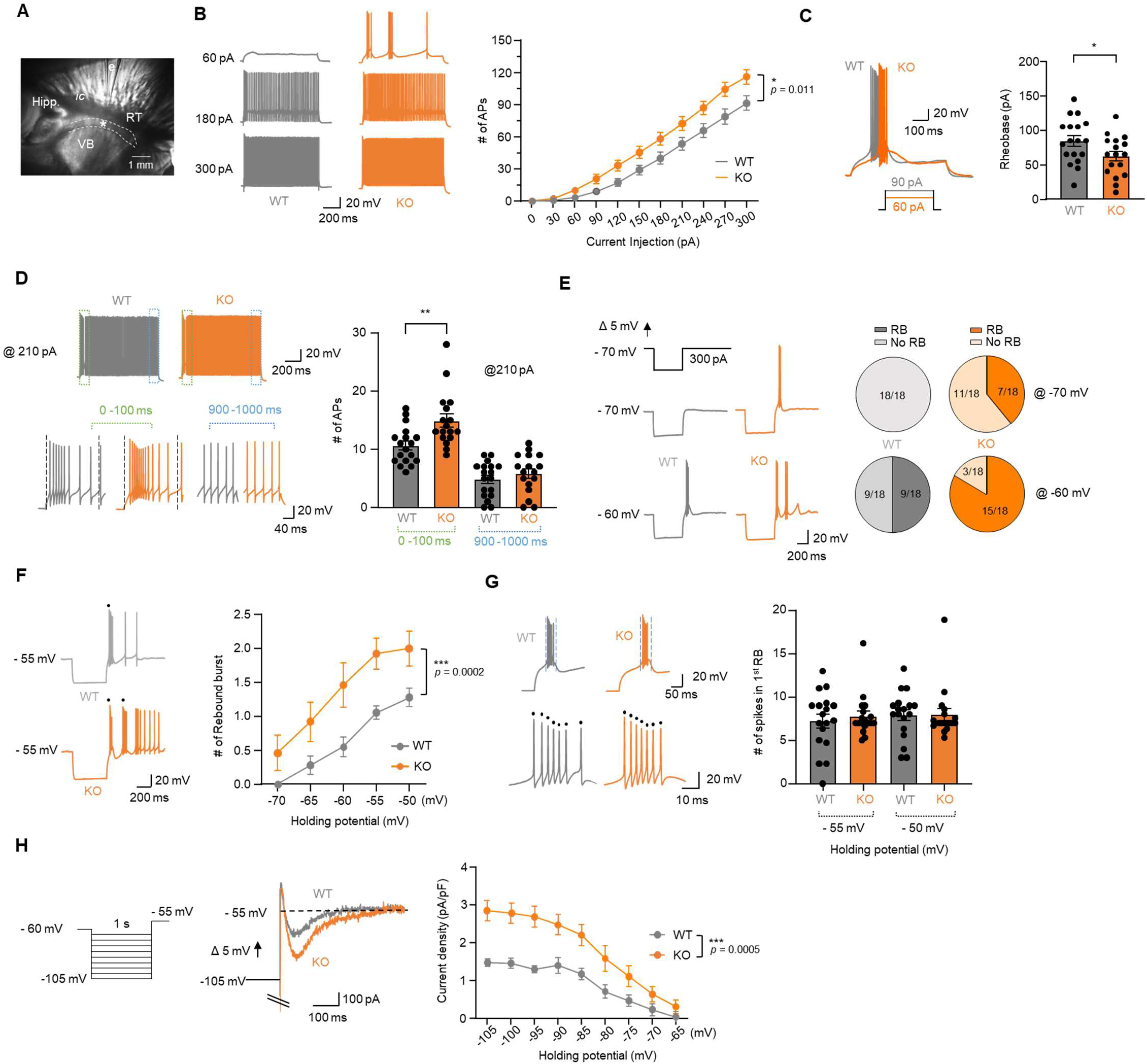
Elevated burst firing in RT underlies increased intrinsic excitability in Cntnap2^−/−^ mice. **(A)** Brightfield image of horizontal thalamic slices with nuclear boundaries in recording chamber, with markings for striatum (Str.), internal capsule (ic), thalamic reticular nucleus (RT), ventrobasal thalamus (VB), hippocampus (Hipp), the glass patch electrode (e), and recording site (*). **(B)** Representative traces showing APs containing bursting and tonic firing elicited by current injection of 60, 180, and 300 pA (Left) and graph showing quantification of the number of APs elicited by 0 – 300 pA (right). **(C)** Representative traces showing APs elicited by injecting rheobase current (left) and quantification of rheobase (right). **(D)** Representative traces showing APs containing an initial burst (within the first 100 ms) followed by tonic firing (measure between 900 – 1000 ms) elicited by injecting 210 pA (Left) in Cntnap2^+/+^ and Cntnap2^−/−^ RT cells. Quantification of the number of burst and tonic APs (right). **(E)** Representative traces showing rebound bursts elicited by injecting hyperpolarizing current (left) and pie charts illustrating the distribution of cells by proportion showing rebound bursts (right). **(F)** Representative traces showing rhythmic oscillatory rebound bursts obtained upon termination of hyperpolarizing current pulses (left), with quantification of the number of rebound bursts as a function of resting membrane potential (right). Black dots indicate single bursts. **(G)** Representative traces showing spikes in the 1^st^ rebound burst (left) and quantification of the number of spikes (right). Black dots each indicate a single spike during the bursts. **(H)** Protocol and representative traces showing low-threshold T-type Ca^2+^currents elicited by membrane hyperpolarization, with subsequent depolarization (left) and summary of current density (pA/pF) of low-threshold T-type Ca^2+^currents (right) (H) n = 15 cells from 3 mice for Cntnap2^+/+^ and n =14 cells from 3 mice for Cntnap2^−/−^. (B-D) n = 18 cells from 4 mice for Cntnap2^+/+^ and n =17 cells from 4 mice for Cntnap2^−/−^. (E-G) n = 18 cells from 4 mice for Cntnap2^+/+^ and n = 18 cells from 4 mice for Cntnap2^−/−^. The statistical tests involved two-way ANOVA (B, F, and H) and an unpaired two-tailed t-test (C and D). Data are represented as mean values ± SEM. *P*-values in figure panels: *p < 0.05, **p < 0.01.

### Altered inhibitory output to TC in Cntnap2^−/−^ mice

Elevation in RT cell excitability in Cntnap2^−/−^ mice is predicted to strengthen RT synaptic outputs. To investigate this, we recorded spontaneous inhibitory postsynaptic currents (sIPSCs) in TC neurons within the VB nuclei. Our results revealed a significant increase in sIPSC frequency, while the amplitude, the rise time and half-width remained unchanged (Fig. 4A), consistent with enhanced RT cell excitability in Cntnap2^−/−^ mice. To further assess synaptic responses evoked by a stimulation of RT neurons in TC neurons, PV- expressing RT neurons were selectively targeted, enabling optogenetic stimulation of RT neurons and their axon terminals (Fig. 4B). Blue light stimulation (1 ms, 470 nm) effectively evoked inhibitory postsynaptic currents (IPSCs) in TC neurons held at –70 mV (Fig. 4C). These IPCSs were confirmed to be mediated by GABA_A_ receptor-dependent chloride currents using two methods: (1) varying holding potentials, between –70 mV, +5 mV (Cl^−^ reversal potential), and +35 mV, and (2) applying SR-95531, a GABA_A_ receptor antagonist, during 5 Hz stimulation (Fig. 4C). After confirming reliable responses to optogenetic stimulation, we assessed presynaptic release dynamics by measuring the paired-pulse ratio (PPR), a marker of presynaptic release probability and observed a significant increase in PPR at 50 ms and 100 ms intervals in Cntnap2^−/−^ mice, indicating alterations in the presynaptic properties of RT inputs onto TC neurons (Fig. 4D). Despite these changes, we observed no differences in synaptic failure rate (Fig. 4E). These findings demonstrate that Cntnap2^−/−^ mice display two synaptic features supporting a presynaptic change in release from RT terminals onto TC cells, an increase in sIPSC frequency and an alteration in release probability that may contribute to the pathophysiology of ASD-related deficits.

**Figure 4.**
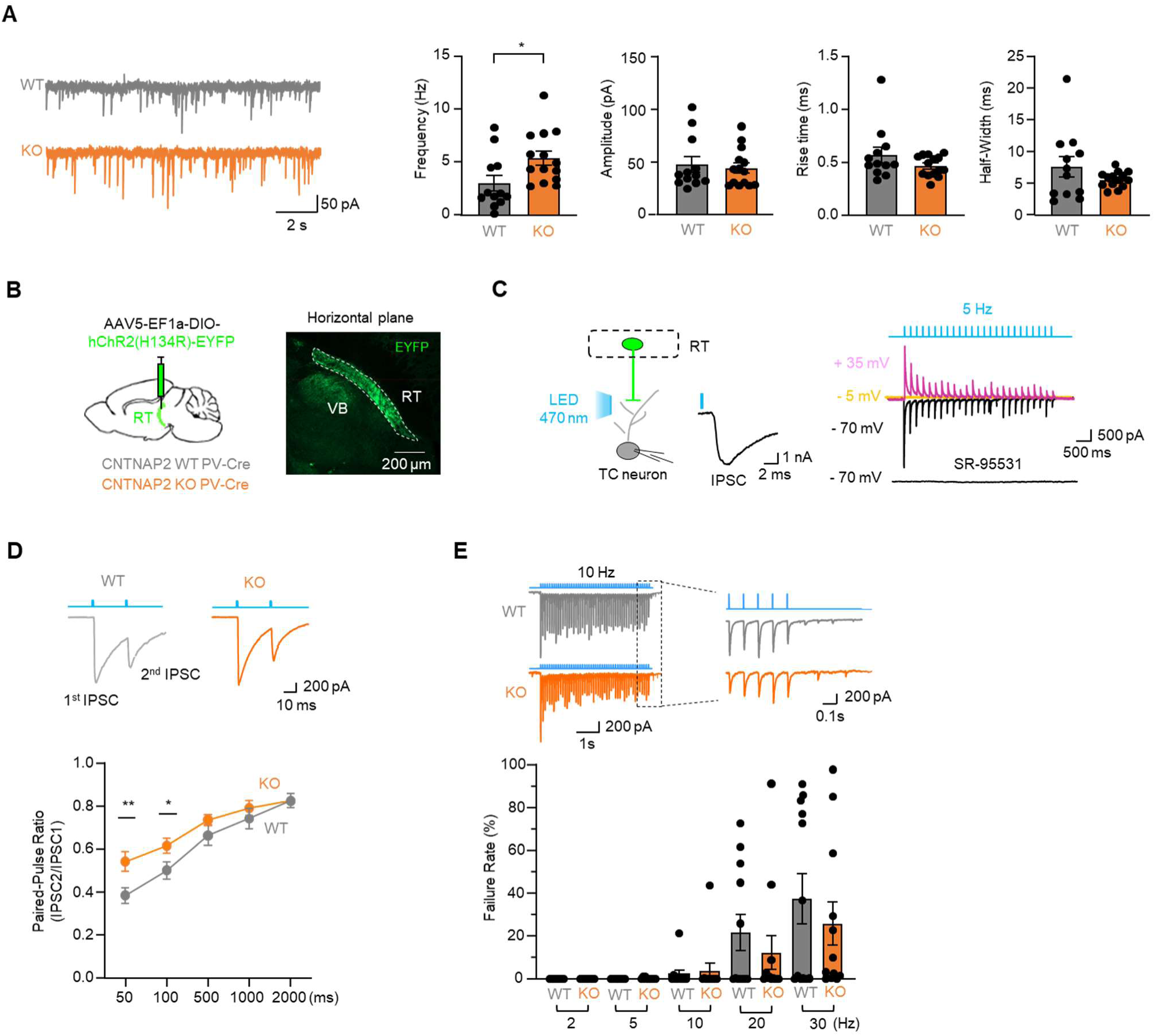
Synaptic inhibitory output from RT->TC is altered in Cntnap2^−/−^ mice. **(A)** Representative traces showing spontaneous IPSCs (left) and quantification of the IPSC frequency (Hz), amplitude (pA), rise time (ms), and half-width (ms) (right). (n = 12 cells from 4 mice for Cntnap2^+/+^ and n = 14 cells from 6 mice for Cntnap2^−/−^). **(B)** Schematic describing stereotaxic injection of AAV5-EF1a-DIO-hChR2(H134R)-EYFP into RT (left) and green fluorescence representing expression of EYFP (right). **(C)** Schematic describing whole-cell patch-clamp recording and a single trace of IPSC elicited by optogenetic stimulation to RT expressing ChR2 (left) and characterization of IPSCs elicited by different membrane holding potentials (−70, −5, and +35 mV) and drug treatment with SR-95531 (right). **(D)** Representative traces showing IPSCs elicited by paired pulses with different time inter-stimulus intervals (50, 100, 500, 1000, and 2000 ms) (top) and quantification of paired-pulse ratio (bottom). (n = 13 cells from 5 mice for Cntnap2^+/+^ and n = 11 cells from 5 mice for Cntnap2^−/−^). **(E)** Representative traces showing IPSCs elicited by 5s-long stimulations at different stimulation frequencies (2, 5, 10, 20, and 30 Hz) (top) and quantification of failure rate (bottom). (n =12 cells from 6 mice for Cntnap2^+/+^ and n =12 cells from 3 mice for Cntnap2^−/−^). The statistical tests involved an unpaired two-tailed t-test (A, D, E). Data are represented as mean values ± SEM. *P*-values in figure panels: *p < 0.05, **p < 0.01.

### Elevated behavior related RT activity in Cntnap2^−/−^ mice

Recent studies using in vivo fiber photometry have shown that RT neurons exhibit spontaneous Ca^2+^ transients in vivo (*64*), reflecting neuronal activity. Based on our observation that intrinsic excitability of RT neurons is increased in Cntnap2^−/−^ mice, we hypothesized that spontaneous in vivo RT population activity would be elevated and exhibit distinct spontaneous responses to external stimuli, including social interactions, objects, and seizures in Cntnap2^−/−^ mice. We confirmed that GCaMP6f was specifically expressed in RT our PV-cre approach (see methods, Fig. 5A). Fiber photometry recordings successfully captured spontaneous calcium transients, quantified using a detection threshold of >3 standard deviations (SD) above baseline (Fig. 5B). Four weeks post-AAV injection, we assessed spontaneous RT activity in response to light and dark conditions, social interactions, inanimate objects, and mild seizures, with 3 - 5 days intervals between stimuli to minimize confounding effects (Fig. 5C). Cntnap2^−/−^ mice exhibited a significant increase in event frequency under illuminated conditions, while no differences were observed in the dark. Cntnap2^−/−^ mice showed an increase in event amplitude (*dF/F*) in the dark without changes in full-width at half-maximum (FWHM) (Fig. 5D). Both Cntnap2^+/+^ and Cntnap2^−/−^ mice displayed increased event frequency during social interaction, while Cntnap2^+/+^ mice also exhibited a reduced event FWHM (Fig. 5E). In response to air puff or tail pinch stimuli, the event amplitude was elevated in Cntnap2^−/−^ mice compared to Cntnap2^+/+^ mice (Fig. S4, A and B). In contrast, no changes in event frequency, peak average, or FWHM were observed in response to inanimate object interaction across both genotypes (Fig. 5F). Finally, consistent with effects of low dose PTZ on seizures (Fig 1A-C) it also increased event frequency in Cntnap2^−/−^ mice, but not in Cntnap2^+/+^ controls (Fig. 5G). These results provide strong evidence that Cntnap2^−/−^ mice exhibit heightened spontaneous activity and stimulus-specific alterations in RT neuron activity, suggesting that increased in vivo population activity may contribute to ASD-related behavioral deficits.

**Figure 5.**
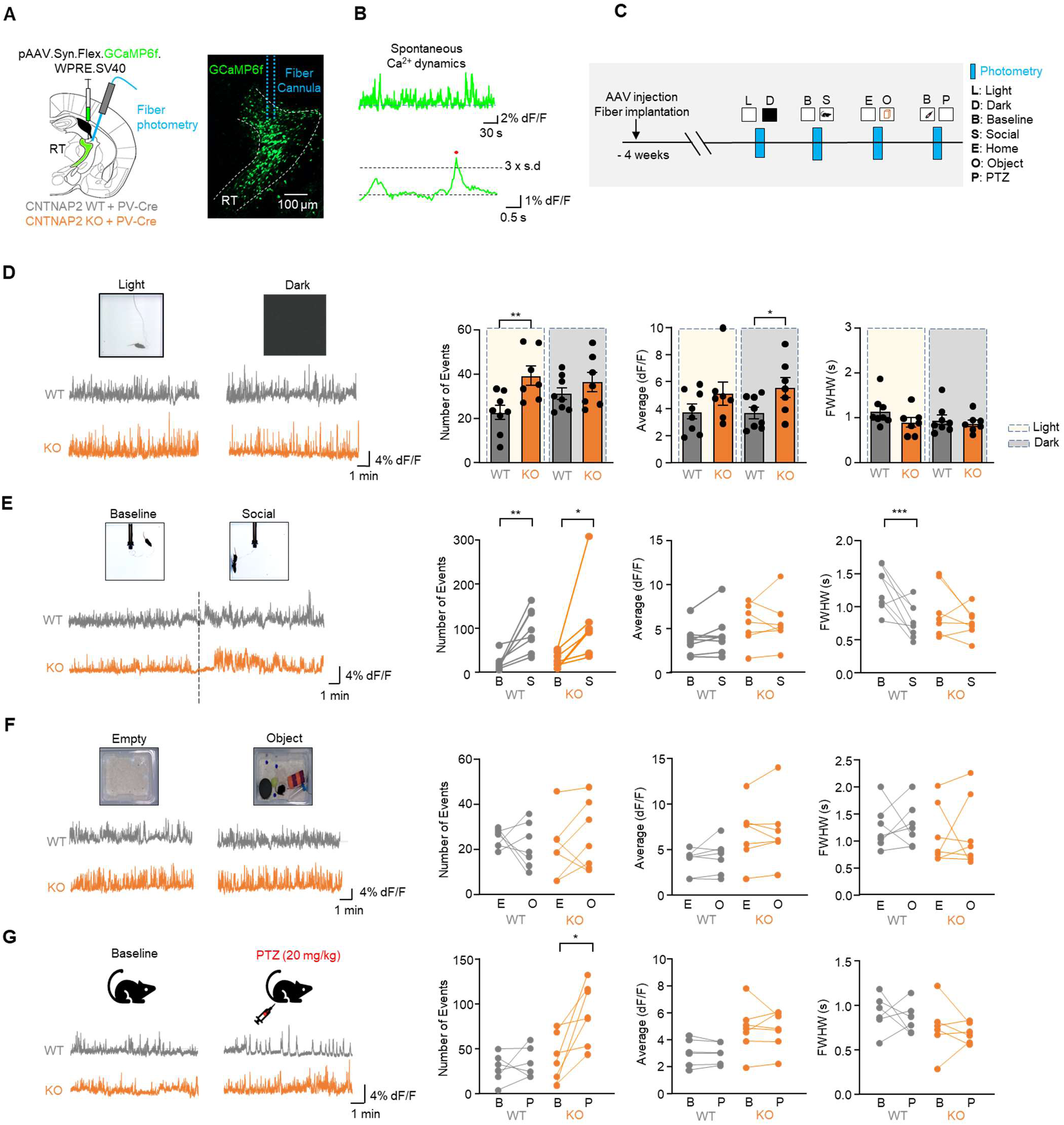
In vivo RT population activity is elevated in Cntnap2^−/−^ mice. **(A)** Schematic describing stereotaxic injection of pAAV.Syn.Flex.GCaMP6f.WPRE.SV40 and fiber cannula into RT regions (left) and green fluorescence representing expression of GCaMP6f and implantation of fiber cannula (right). **(B)** Representative traces showing spontaneous Ca^2+^ dynamics (top) and detection criteria (bottom). **(C)** Schematic illustrating experimental conditions and strategy to measure spontaneous Ca^2+^ dynamics in response to a variety of stimuli. **(D)** Representative images showing experimental conditions (top, left) and traces of spontaneous Ca^2+^ dynamics during light and dark conditions (bottom, left) and quantification of number of events, average (dF/F), and FWHM in Ca^2+^ dynamics (right). (n = 8 mice for Cntnap2^+/+^ and n = 7 mice for Cntnap2^−/−^). **(E)** Representative images showing experiment (top, left) and spontaneous Ca^2+^ levels during baseline and social condition (bottom, left) and quantification of events (right). (n = 8 mice for Cntnap2^+/+^ and n = 7 mice for Cntnap2^−/−^). **(F)** Representative images showing experiment (top, left) and traces Ca^2+^ levels during empty and object conditions (bottom, left) and quantification of events (right). (n = 7 mice for Cntnap2^+/+^ and n = 7 mice for Cntnap2^−/−^). **(G)** Schematic of experiment (top, left) and spontaneous Ca^2+^ levels during baseline and PTZ condition (bottom, left) and quantification of events (right). (n = 6 mice for Cntnap2^+/+^ and n = 6 mice for Cntnap2^−/−^). The statistical tests involved two-way ANOVA with Tukey’s test (D) and a paired two-tailed t-test (E-G). Data are represented as mean values ± SEM. *P*-values in figure panels: *p < 0.05, **p < 0.01, ***p < 0.001.

### Acute Systemic inhibition of T-type calcium channel restores behavioral deficits in Cntnap2^−/−^ mice

Given the correlative relationship between behavioral deficits and elevated T current dependent burst firing in RT of Cntnap2^−/−^ mice (Fig. 3, F and G), we hypothesized that inhibiting T-type calcium channels could rescue behavioral deficits observed in Cntnap2^−/−^ mice. Before conducting behavioral rescue experiments, we validated the effects of Z944, a novel, orally bioavailable, and selective T-type calcium channel blocker (*65*), on RT activity using fiber photometry in vivo and whole-cell patch clamp in vitro. Systemic administration of Z944 (10 mg/kg) significantly reduced the number of calcium transients without affecting amplitude or FWHM (Fig. S5A). Bath application of Z944 at 0.1 µM and 1 µM Z944 significantly increased burst rheobase (Fig. S5B) and reduced the number of APs/burst (Fig. S5C) in a dose-dependent manner. These data confirm effective pharmacological inhibition of in vivo RT population activity and intrinsic excitability in RT neurons. Behavioral tests with Z944 were performed at 3–5 days between vehicle controls and drug treatments to allow recovery and avoid confounding effects of repeated behavioral tests (Fig. 6A). During the 10-minute habituation phase, Cntnap2^−/−^ mice displayed hyperlocomotion compared to Cntnap2^+/+^ mice, which was normalized by Z944 to values near those seen in Cntnap2^+/+^ mice (Fig. S7A). Similarly, in the subsequent 20- minute test session, Cntnap2^−/−^ mice continued to travel further than Cntnap2^+/+^ mice, and this difference was attenuated by Z944, reducing locomotion to Cntnap2^+/+^ levels (Fig. 6B). The distance traveled by Cntnap2^+/+^ mice was unaffected by either treatment (Fig. 6B). In the 3-chamber social preference test, vehicle treated Cntnap2^+/+^ mice spent significantly more time in the chamber containing the social partner, demonstrating a normal social preference (Fig. 6C). In contrast, Cntnap2^−/−^ mice treated with vehicle exhibited no such preference, replicating our results in untreated mice (Fig. 1E), spending equal time exploring both chambers (Fig. 6C). Notably, Z944 treatment rescued social behavior in Cntnap2^−/−^ mice, as shown by increased time spent in the chamber containing the social partner (Fig. 6C). Manually scoring of sniffing interaction time gave complementary results (Fig. S7B). Z944 normalized the elevated grooming time observed in Cntnap2^−/−^ mice to levels seen in Cntnap2^+/+^ mice treated with either vehicle or Z944 (Fig. 6D) with no changes in the number of grooming events (Fig. 6E) or digging time (Fig. 6F). Cntnap2^−/−^ mice displayed reduced time interacting in the reciprocal interaction test compared to Cntnap2^+/+^ mice. Interestingly, Z944 reduced interaction time in Cntnap2^+/+^ mice and restored the interaction time in Cntnap2^−/−^ mice (Fig. 6G). These rescue experiments using Z944 demonstrate that systemic inhibition of T-type calcium channels effectively mitigated hyper-locomotor activity, restored social behaviors, and reduced repetitive behaviors in Cntnap2^−/−^ mice, highlighting its therapeutic potential for treating behavioral deficits associated with ASD.

**Figure 6.**
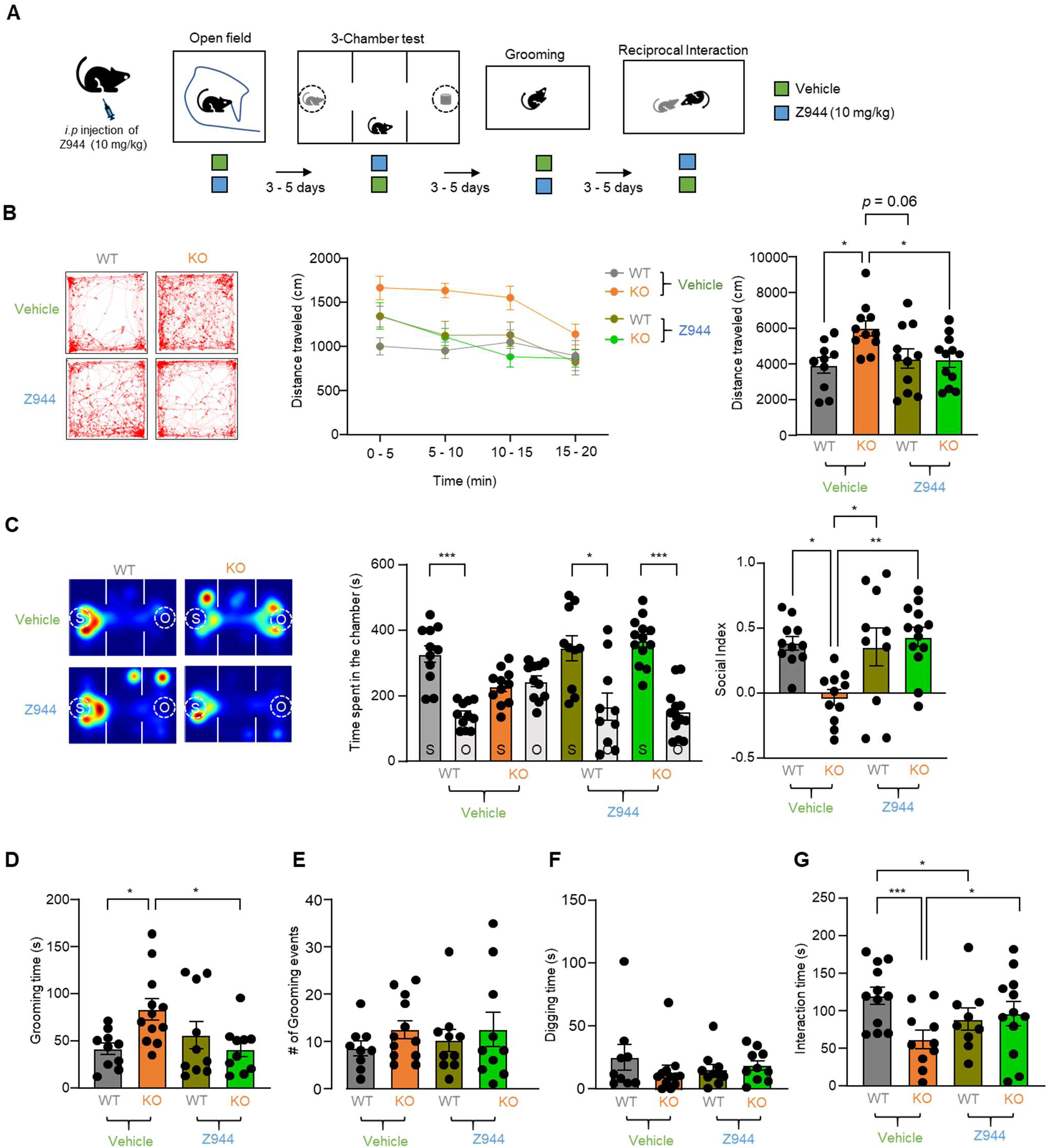
Pharmacological inhibition of T-type calcium channels ameliorates ASD- related behaviors. **(A)** Schematic of experiment for evaluating the rescue effect of Z944 (10 mg/kg), a T-type calcium channel inhibitor, on behavioral phenotypes. **(B)** Representative open field movement traces (left), and summary of distance traveled in 5-min intervals (middle), along with quantification of total distance traveled over 20 mins (right) in Cntnap2^+/+^ and Cntnap2^−/−^ mice treated with vehicle or Z944. (n = 10 mice for Cntnap2^+/+^-Veh, n = 11 for Cntnap2^−/−^-Veh, n = 11 for Cntnap2^+/+^-Z944, n = 12 for Cntnap2^−/−^-Z944). **(C)** Representative heatmaps showing location density across time during social preference tests (left), quantification of time spent in each chamber (middle) and the social index (right) in Cntnap2^+/+^ and Cntnap2^−/−^ mice treated with vehicle or Z944. (n = 11 mice for Cntnap2^+/+^ -Veh, n = 11 for Cntnap2^−/−^-Veh, n = 10 for Cntnap2^+/+^ -Z944, n = 13 for Cntnap2^−/−^-Z944). **(D-F)** Quantification of grooming time (D), the number of grooming events (E), and digging time (F) in Cntnap2^+/+^ and Cntnap2^−/−^ mice treated with vehicle or Z944. (n = 10 mice for Cntnap2^+/+^ -Veh, n = 12 for Cntnap2^−/−^-Veh, n = 10 for Cntnap2^+/+^ - Z944, n = 10 for Cntnap2^−/−^-Z944). **(G)** Quantification of interaction time in Cntnap2^+/+^ and Cntnap2^−/−^ mice treated with vehicle or Z944. (n = 12 mice for Cntnap2^+/+^ -Veh, n = 10 for Cntnap2^−/−^-Veh, n = 9 for Cntnap2^+/+^ -Z944, n = 12 for Cntnap2^−/−^-Z944). The statistical tests involved two-way ANOVA with Tukey’s test (B-D). Data are represented as mean values ± SEM. *P*-values in figure panels: *p < 0.05, **p < 0.01, ***p < 0.001.

### Acute DREADD-mediated inhibition of RT restores behavioral deficits in Cntnap2^−/−^ mice

Lastly, to test whether broadly suppressing RT activity might mitigate behavioral deficits in Cntnap2^−/−^ mice, we selectively expressed hM4Di-mCherry in RT without significant spillover (see methods, Fig. 7A). In vitro experiments with whole-cell patch clamp recording in slices confirmed that hM4di was expressed in RT cells, and that C21 decreased electrical excitability. Administration of C21 (5 µM) in the bath hyperpolarized membrane potentials and increased rheobase (Fig. 7B). Once confirmation of functional hM4Di expression of RT cells was complete, behavioral tests were conducted at intervals of 3-5 days to allow recovery and minimize potential confounding effects (Fig. 7C). In the open field test, during the 10-minute habituation phase, Cntnap2^−/−^ mice displayed increased locomotor activity compared to Cntnap2^+/+^ mice treated with vehicle. C21 injection failed to rescue this hyperactivity (Fig. S7C). During the 20-minute test session, however, Cntnap2^−/−^ mice hyperactivity was significantly attenuated by C21, reducing the distance traveled to Cntnap2^+/+^ levels, while the locomotor activity in Cntnap2^+/+^ remained unaffected by either treatment. (Fig. 7D). In the 3-chamber social preference test, vehicle-treated Cntnap2^−/−^ mice exhibited no preference for the social chamber while C21 injection restored social preference in Cntnap2^−/−^ mice (Fig. 7E). In the grooming test, C21 injection significantly increased grooming time in Cntnap2^+/+^ mice (Fig. 7F) and the number of grooming events (Fig. 7G), suggesting a role for RT in regulating repetitive behaviors. Notably, C21 injection rescued elevated grooming time and the number of grooming events in Cntnap2^−/−^ mice, similar to Cntnap2^+/+^ levels (Fig. 7, F and G). However, there was no effect on digging time (Fig. 7H). C21 failed to rescue the reduced interaction time in Cntnap2^−/−^ mice (Fig. 7I). Given that reduction of RT activity with hM4D/C21 rescued ASD relate behaviors in Cntnap2^−/−^ mice, we tested whether increased RT activation with hM3D/C21 would evoke ASD type behaviors. Specific expression of hM3Dq-mCherry in RT was confirmed with minimal or no spillover into neighboring regions, such as the zona incerta (Fig. S6A). C21 (5 µM) decreased rheobase currents and depolarized membrane potentials in vitro (Fig. S6B). Behaviorally, Cntnap2^+/+^ mice treated with vehicle showed social preference while C21 activation of RT neurons expressing hM3Dq reduced social preference, with mice spending equal time in either chamber and exhibiting a reduced social index (Fig. S6C). C21 injection increased grooming time, suggesting that elevated RT activity can induce ASD-related behavioral deficits similar to those observed in Cntnap2^−/−^ mice (Fig. S6D). Our behavioral data using DREADDs highlight the critical role of RT activity in mediating ASD-related behaviors, including hyper-locomotor activity, repetitive behaviors, and social deficits in Cntnap2^−/−^ mice, providing insights into its potential target for alleviating ASD-related behaviors.

**Figure.7.**
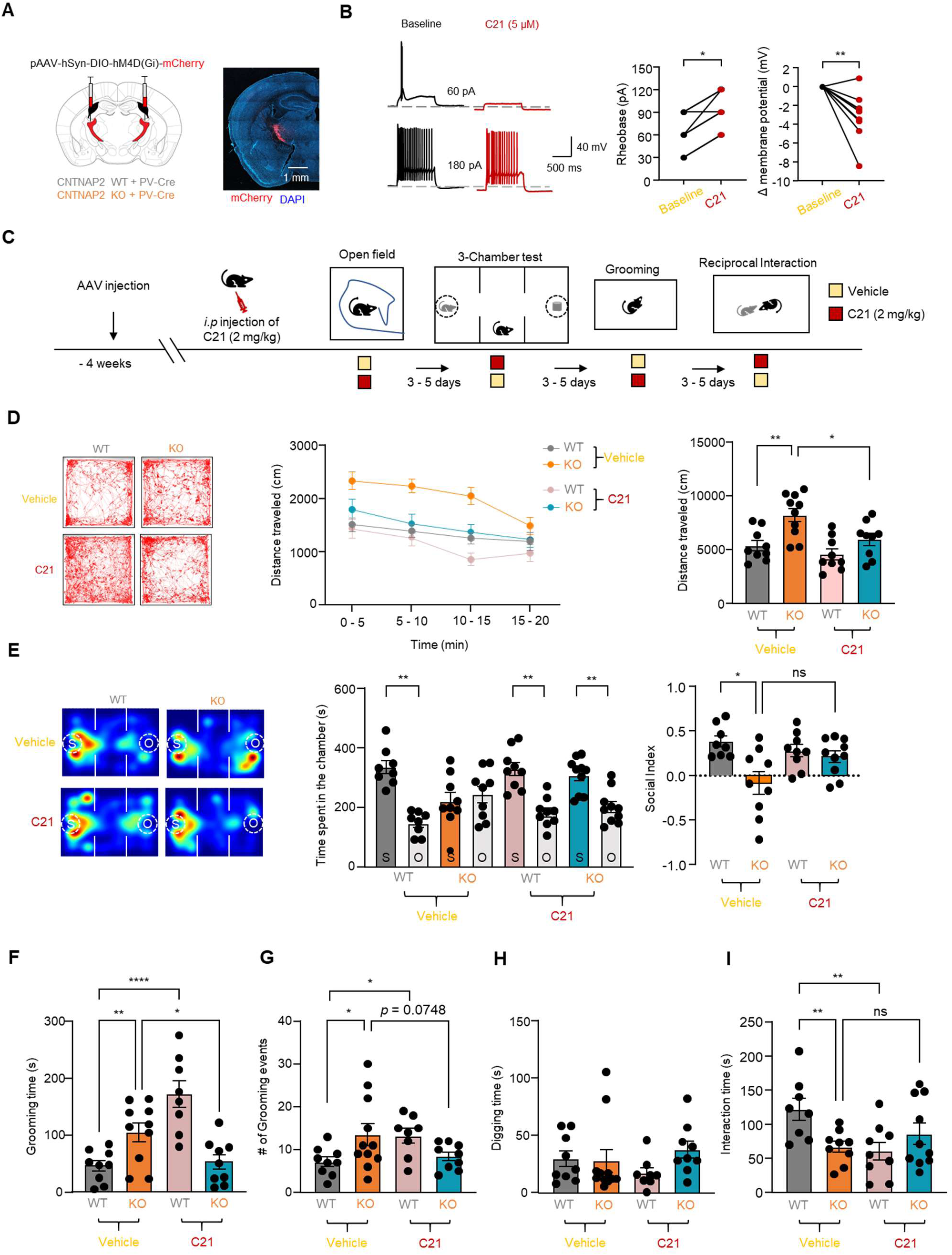
Chemogenetic inhibition of RT ameliorates ASD-related behaviors. **(A)** Schematic describing stereotaxic injection of pAAV-hSyn-DIO-hM4D(Gi)-mCherry virus into RT (left) and red fluorescence representing expression of mCherry (right). **(B)** Representative traces showing APs (left) elicited by rheobase currents and quantification of C21- induced changes in membrane potential and rheobase (right). (n = 9 cells from 4 mice). **(C)** Experimental strategy for evaluating the rescue effect of chemogenetic inhibition on behavioral phenotypes. **(D)** Representative open field movement traces (left), distance traveled in 5-min intervals (middle), and quantification of total distance traveled over 20 mins (right) in Cntnap2^+/+^ and Cntnap2^−/−^ mice treated with vehicle or C21. (n = 9 mice for Cntnap2^+/+^-Veh, n = 10 for Cntnap2^−/−^-Veh, n = 9 for Cntnap2^+/+^-C21, n = 9 for Cntnap2^−/−^-C21). **(E)** Representative heatmaps showing location data during social preference tests (left), time spent in the chamber (middle) and the social index (right) in Cntnap2^+/+^ and Cntnap2^−/−^ mice treated with vehicle or C21. (n = 8 mice for Cntnap2^+/+^- Veh, n = 9 for Cntnap2^−/−^-Veh, n = 9 for Cntnap2^+/+^-C21, n = 10 for Cntnap2^−/−^-C21). **(F- H)** Quantification of grooming time (F), the number of grooming events (G), and digging time (H) in Cntnap2^+/+^ and Cntnap2^−/−^ mice treated with vehicle or C21. (n = 9 mice for Cntnap2^+/+^-Veh, n = 7 for Cntnap2^−/−^-Veh, n = 8 for Cntnap2^+/+^-C21, n = 9 for Cntnap2^−/−^- C21). **(I)** Quantification of interaction time in Cntnap2^+/+^ and Cntnap2^−/−^ mice treated with vehicle or C21. (n = 8 mice for Cntnap2^+/+^-Veh, n = 9 for Cntnap2^−/−^-Veh, n = 9 for Cntnap2^+/+^-C21, n = 11 for Cntnap2^−/−^-C21). The statistical tests involved a paired two-tailed t-test (B) and two-way ANOVA with Tukey’s test (D-F). Data are represented as mean values ± SEM. *P*-values in figure panels: *p < 0.05, **p < 0.01, ***p < 0.001, ns, not significant.

## Discussion

In the present study, we provide comprehensive evidence that RT neurons, a critical inhibitory component of thalamocortical circuits, are hyperexcitable in the Cntnap2^−/−^ model of ASD. We find an increase in bursting activity of RT neurons, a disruption of their synaptic output to TC neurons, and elevated spontaneous RT activity in vivo. Importantly, pharmacological and chemogenetic suppression of RT excitability ameliorates key ASD- related behavioral abnormalities, such as hyper-locomotive activity, impaired social preference, and repetitive behaviors. These findings establish RT hyperexcitability as a key contributor to ASD pathophysiology and highlight its potential as a therapeutic target.

Over the past decades, individuals with ASD and animal models have been characterized by common co-occurring symptoms such as heightened tactile sensitivity (*66*, *67*), attention-deficit/hyperactivity disorder (ADHD) (*27*), sleep disturbances(*12*, *13*, *40*), and epilepsy(*1*, *14*, *45*, *68*). Previous studies in Cntnap2^−/−^ mice have also reported increased startle response (*69*), abnormal motor behavior (*1*), disrupted sleep-wake cycle (*40*), degraded tactile sensation (*4*), and spontaneous seizure-like spike discharge (*40*). Our in vivo EEG and behavioral analysis in Cntnap2^−/−^ mice confirmed increased seizure susceptibility and heightened locomotor activity, both of which are associated with epilepsy, alongside core ASD symptoms. This underscores the need to elucidate shared neural mechanisms underlying these comorbidities to optimize therapeutic strategies. Given the well-established link between ASD and epilepsy, identifying neural circuits that contribute to both conditions is crucial. The thalamus, a central hub for sensory processing and cognitive functions necessary for social communication, remains an underexplored yet promising candidate in understanding ASD-related pathophysiology. In human ASD patients, atypical thalamocortical connectivity has been reported to contribute to ASD symptoms (*18*, *70*). SHANK3 KO mice, a widely studied ASD model, were reported to exhibit increased thalamocortical spike and burst rates, accompanied by ASD-like behavioral phenotype (*19*). Similarly, our findings demonstrate heightened spontaneous and evoked synchronized oscillatory activity within thalamocortical networks in Cntnap2^−/−^ mice. Together, these observations suggest that thalamocortical network dysfunction, driven by altered oscillatory dynamics, might play a pivotal role in the pathophysiology of ASD and offers a promising target for therapeutic intervention.

Thalamic oscillations rely on intact reciprocal connection between RT and TC neurons. As a potent driver of oscillatory activity, RT exhibits intrinsic oscillations (*71*, *72*) and exerts strong synaptic inhibition on TC cells. Upon RT activation, TC neurons become hyperpolarized, triggering rebound bursts that re-excite RT. This interaction initiates a cycle of oscillatory activity, often described as a “ping-pong” dynamic between RT and TC (*73*, *74*). Disruptions in RT activity, whether through activation or inhibition, could induce absence seizures, characterized by hypersynchronous activity (*28*, *58*). Notably, increased RT bursting was reported in a mouse model of Dravet syndrome (*51*), a severe form of epilepsy, supporting our conclusion that heightened bursting activity in RT neurons contributes to the elevated seizure susceptibility and behavioral deficits observed in Cntnap2^−/−^ mice. In the context of ASD, a recent study highlights RT activity during social interactions and its role in encoding information for recognizing familiar conspecifics (*75*), supporting a role for RT in social behaviors. Consistent with this, our study demonstrates that pharmacological and chemogenetic inhibition of hyperexcitability and burst firing in RT neurons of Cntnap2^−/−^ mice ameliorates ASD-related deficits, whereas enhanced RT activity impairs social preference and grooming behavior. Interestingly, both RT activation and inhibition lead to elevated grooming behaviors, suggesting a critical role of balanced RT activity in maintaining homeostasis.

RT neurons have functional synapses with various brain regions, including various nuclei in dorsal thalamus, such as the TC cells in ventrobasal complex (VB), and the lateral habenula (LHb), and roles associated with each connection have been identified. Reduction in spontaneous RT activity and RT-to-TC projections are associated with chronic sleep disruption (CSD)-induced hyperalgesia (*64*) and acute manipulation of the somatosensory RT-to-lateral habenula pathway induces or prevents depressive-like behaviors, indicating that RT neurons are linked to pain and emotional processing (*39*). Our data that altered presynaptic release probability and spontaneous output of RT neurons to TC also suggest that disruption in synaptic plasticity at the RT-TC synapse might contribute to ASD-related behaviors in Cntnap2^−/−^ mice. Future research should address the specific roles of RT neurons and their connection with other brain regions in regulating fundamental processes such as emotional, cognitive, and sensory processes that influence social behavior and locomotive activity. A deeper understating of these mechanisms will provide valuable insights into RT’s contributions to ASD physiology and its potential as a therapeutic target.

RT neurons comprises a relatively homogeneous population of GABAergic neurons (*76*), yet there is heterogeneity in molecular and electrophysiological properties, connectivity, and functional roles (*22*, *63*). Most RT neurons express PV, and have distinct firing properties and specific projections to TC neurons, critical for modulating somatosensory behavior and seizure activity (*20*). Notably, reduced PV neuron counts and decreased PV expression have been reported in postmortem brains of individuals with ASD (*5*, *60*, *77*). Similarly, Cntnap2^−/−^ show a reduction in PV+ cell counts, decreased spontaneous mIPSC frequency, and diminished feedforward eIPCSs in striatal projection neurons (SPNs) (*2*). These findings support the “PV hypothesis” (*78*), which posits that reduced PV levels impair intrinsic excitability, synaptic function, network properties, all of which are causally linked to the etiology of ASD. Given the high density of PV-expressing interneurons in RT (*57*), RT neurons represent a promising target for further investigation into the mechanisms underlying ASD. In our in vitro intracellular recordings, slices from Cntnap2^−/−^ mice exhibited increased rebound bursts and a hyperpolarizing shift for burst activation threshold in RT neurons, implicating enhanced burst firing in ASD-related deficits. This hypothesis is further supported by recent findings, including (1) reduced bursting firing in RT neurons of Ptchd1^Y/-^ mice displaying attention deficits and hyperactivity (*27*) and (2) transformations of burst patterns in RT neurons during locomotion (*79*). Mechanistically, T-type calcium channels play a crucial role in regulating bursting activity in RT neurons, driving synchronized rhythmicity within the thalamic circuits (*80*). Disruptions in this low-voltage activated calcium influx can lead to aberrant bursting, contributing to conditions such as epilepsy and impairments in sensory and cognitive processing. A recent study in Gclm-KO mice, a schizophrenia model, revealed that decreased Cav3.3 expression and hypofunction of T-type calcium channels in RT neurons lead to a hyperpolarizing shift in the threshold for repetitive bursting, thereby altering firing properties (*37*). In addition, a genome-wide association study (GWAS) has identified single nucleotide polymorphisms (SNPs) in the Cav3.3 gene as risk factor for autism and neuropsychiatric disorders (*81*). Given that the deletion of Cav3.3 channels, which are highly enriched in RT (*82*), reduces T-type calcium currents in RT by approximately 80% and abolishes rebound bursts (*83*), we anticipated that enhanced Cav3.3 function could elevate T-type calcium currents in RT neurons of Cntnap2^−/−^, which potentially contributes to ASD-related behaviors.

Z944, a glycinamide compound, has been established as an effective blocker of bursting firing and T-type calcium currents in RT neurons (*65*). Its therapeutic potential has been extensively explored in animal models, demonstrating efficacy in alleviating pain (*84*), absence seizures (*65*), and deficits in learning and locomotor function (*85*). Notably, systemic administration of Z944 reduces introductory and aggressive behaviors associated with social communication in the Genetic Absence Epilepsy Rats from Strasbourg (GAERS) (*86*) and we find that a single injection of Z944 is effective to restore behavioral deficits in Cntnap2^−/−^ mice, strongly suggesting that systemic inhibition of T-type calcium channel might be a promising approach for addressing ASD-related behavioral deficits. We cannot at this point exclude the possibility that the effects of Z944 on behavior were restricted to RT, with potential contributions from other brain regions with prominent Cav3 expression. To better understand the distinct contribution of T-type calcium channel in RT neurons, future studies might target reduced functional expression of RT Cav3 channels using siRNA or short hairpin RNA (shRNA) or CRISPR-based gene editing. In any case, a key advantage of Z944 is its oral bioavailability, enabling safe and sustained administration at different stages of development, such that early interventions would be possible. A study on VPM hyperactivity due to HCN2 dysfunction demonstrated that early developmental treatments could produce lasting effects (*19*). In that study, HCN activation through lamotrigine (LTG) treatment resulted in long-lasting rescue effects that persisted beyond its acute benefits. Although our study did not explore changes in RT bursting activity during postnatal development or the consequent behavioral abnormalities that manifest in adulthood, further studies should address the onset of RT dysfunction and explore the potential for early intervention with oral administration of Z944 to restore normal function over time.

While the pharmacological approach with Z944 did not direct support a role of RT T channels, per se, in ASD behaviors, the results of excitatory and inhibitory DREADDs do provide complementary support for an RT role, as the PV-cre and viral approach we used specifically targeted RT cells. In general, DREADD approaches are providing novel therapeutic possibilities. For example, targeting RT activity was shown to rescue deficits in an Alzheimer’s disease (AD) model (*87*) and improved prepulse inhibition (PPI) of the acoustic startle response, which has been linked to neuropsychiatric disorders (*69*), highlighting the therapeutic potential of selective RT manipulation for achieving pharmacological benefits. Similar to the effects of Z944, in our hands when RT activity is generally suppressed by an inhibitory DREADD approach, ASD pathological behaviors were rescued in Cntnap2^−/−^ mice. Finally, in support of a causative role of RT in driving ASD related behaviors, excitatory DREADDs designed to increase RT activity in Cntnap2^+/+^ mice initiated ASD-related behaviors such as deficits in social preference and grooming.

Overall, this study identifies elevated RT burst firing and aberrant thalamic oscillatory dynamics in Cntnap2^−/−^ mice as a key driver of ASD-related behavioral deficits. If this is a common mechanism of ASD-circuit pathology arising from a variety of genetic causes, then compounds such as Z944, or subtype specific T-type calcium channel antagonists that would target the Cav3.2 and Cav3.3 expressed in RT neurons, might be an effective therapeutic strategy. Furthermore, future research should focus on elucidating RT’s roles in sensory, emotional, and sleep regulation to optimize therapeutic strategies in the context of ASD.

## Materials and Methods

### Animals

Experimental Cntnap2^+/+^ (wild-type, WT) and Cntnap2^−/−^ (homozygous knockout, KO) mice were obtained from heterozygous crosses of Cntnap2^+/-^ (heterozygous, HET) mice and used for electroencephalogram (EEG) recording, behavioral tests, electrophysiology in acute brain slices, and pharmacology experiments. For fiber photometry, optogenetic, and chemogenetic experiments, Cntnap2^+/+^ x PV-Cre and Cntnap2^−/−^ x PV-Cre mice were generated by crossing male and female Cntnap2^+/-^ (HET) x PV-Cre mice, derived from crossing Cntnap2^−/−^ mice x PV-Cre mice (Jax Stock#: 008069). For optogenetically induced oscillation experiments, Cntnap2^+/+^ mice x Ai32/Ntsr1-Cre and Cntnap2^−/−^ mice x Ai32/Ntsr1-Cre mice were produced by crossing male and female Cntnap2^+/-^ (HET) x Ai32/Ntsr1-Cre, which were generated from Ai32 (Jax Stock#: 012569) with Ntsr1-cre mice (gift from University of California, Davis). All mice were maintained on a reverse 12-hour dark/light cycle, with experiments conducted during their active (dark) phase. Male mice aged 8–14 weeks were used in all experiments except for oscillation recordings, in which equal numbers of males and females were used. Food and water were provided ad libitum. All experimental procedures were approved by the Stanford Administrative Panel on Laboratory Animal Care (APLAC, Protocol #12363) and conducted in accordance with National Institutes of Health (NIH) guidelines.

### Virus Injection and Stereotaxic Surgery

Mice were anesthetized with continuous isoflurane (3% for induction, 0.8%-1.5% for maintenance) and placed in a stereotaxic rig for surgery. A heating pad was used during anesthesia induction and postoperatively, and ophthalmic ointment was applied to protect the eyes throughout the procedure. Stereotaxic coordinates for injections/implantations were as follows: RT (virus injection) - AP: - 1.10 mm, ML: ± 2.00 mm, DV: 3.10 mm. Viral infusions were performed using a 10-minute injection (50 nL/min), with a total volume of 350 nL for ex vivo electrophysiology experiments at the RT-TC synapse and chemogenetic modulation. Cntnap2^+/+^ mice x PV-Cre and Cntnap2^−/−^ mice x PV-Cre mice received bilateral injections of AAV2-DIO-hChR2-eYFP, AAV2-hSyn-DIO-hM4D(Gi)-mCherry, or AAV2-hSyn-DIO-hM3D(Gq)-mCherry into the RT, obtained from Addgene (plasmid#26973, plasmid#44362, and plasmid#44361). After injection, needles were held in place for 10 minutes before withdrawal to minimize backflow. Mice were allowed to recover for 3-4 weeks before undergoing experiments, including fiber photometry, behavioral assays, electrophysiology, or histological analysis.

### Electroencephalogram (EEG)

Cntnap2^+/+^ and Cntnap2^−/−^ mice, aged 2–4 months, were used for EEG recordings. Mice were anesthetized with isoflurane during the EEG electrode implantation procedure. Under anesthesia, the skull was exposed, and small holes were drilled bilaterally at specific stereotaxic coordinates (AP −1.0 mm, ML ± 2.0 mm) for the somatosensory cortex to accommodate electrode placement. Sterilized gold pins were secured to the skull at the designated recording sites using adhesive, with a reference electrode positioned in the cerebellum. Following electrode placement, the mice were allowed to recover in their home cages for at least three days to ensure stable electrode integration. EEG recordings were conducted in a quiet, controlled environment during the (dark) active phase. Mice were connected to an Open Ephys/Intan headstage recording system via lightweight, flexible cables for data acquisition. EEG signals were amplified and filtered at 2 kHz, with continuous recordings performed for 10 min following intraperitoneal (i.p) injection of pentylenetetrazole (PTZ; Sigma; 20 mg/kg). EEG data were analyzed offline using custom MATLAB scripts to manually process the recorded signals.

### Behavioral tests

Behavioral tests were conducted using male mice, aged 8–12 weeks, because Cntnap2^−/−^ female mice show no significant behavioral deficits (*43*). Tests were performed during light-off periods. Subject mice were given at least 2 days of rest between different tests to minimize stress or carryover effects. Behavioral data were recorded using a video camera positioned above the testing area. Analysis was performed manually or using custom MATLAB scripts designed for the specific behavioral paradigms.

### Open-field test

A subject mouse was placed into the center of the open-field box (white acryl chamber, 50 x 50 x 50 cm) and recorded for 10 min for Cntnap2^+/+^ and Cntnap2^−/−^ mice comparisons, or 30 min for Z944 and DREADD experiments. The chamber was illuminated at ∼ 100 lux. Mouse movements were tracked and analyzed using customized Python scripts.

### Three-chamber test

Subject mice were isolated for 24 hours prior to testing. The three-chambered apparatus (white acrylic chamber, 40 × 20 × 25 cm) consisted of left, center, and right chambers arranged in a row, with two entrances connecting the center chamber to the side chambers. The test consisted of two 10-minute phases. In the first phase the subject mouse was introduced into the center chamber and allowed to freely explore all three chambers with empty wire cages positioned in the left and right chambers. In the second phase, one wire cage was replaced with a novel object (O), while another cage contained a stranger mouse (S) which were C57BL/6J strain and sex- and age-matched to the subject mouse. During the interval between phases, the subject mouse was gently guided to the center chamber, and the entrances to the side chambers were temporarily blocked by barriers and then the barriers were removed to let the subject mouse move freely in the arena. The center chamber was illuminated at ∼100 lux. Time spent in the chambers containing the stranger mouse or object was recorded and analyzed using custom Python scripts. The sociality index (*SI*) was calculated as [*T*_mouse_−*T*_object_]/[*T*_mouse_+*T*_object_], where *T*_mouse_ and *T*_object_ represent the time spent interacting with the stranger mouse and object, respectively. To further obtain detailed information about social interaction, time spent in direct physical contact with wire cages on both sides was manually scored in a blind manner. Between each measurement, the apparatus was cleaned with 75% ethanol to minimize residual olfactory cues.

### Grooming behavior

A subject mouse was placed in a new home cage containing fresh bedding and allowed to freely explore for 20 minutes. The center of the cage was illuminated at approximately 100 lux. Time spent digging and self-grooming during the final 10 minutes was recorded manually in a blinded manner. Digging was defined as the mouse using its head or forelimbs to displace bedding. Self-grooming was defined as the mouse stroking or scratching its face or body, or licking its body.

### Reciprocal Interaction

A subject mouse, aged 8–12 weeks, was placed in a chamber (40 × 20 × 25 cm) and allowed to habituate for 10 minutes. Following habituation, a novel conspecific mouse, matched for genotype, age, sex, and/or treatment, was introduced into a neutral area of the chamber. The time spent in social interactions, including nose-to-anogenital sniffing and nose-to-nose sniffing, was recorded manually by an observer blinded to the experimental conditions of the test mouse.

### Thalamic Slice preparation

Mice were anesthetized with isoflurane and transcardially perfused with ice-cold sucrose-based cutting solution containing (in mM) 234 sucrose, 11 glucose, 26 NaHCO_3_, 2.5 KCl, 1.25 NaH_2_PO_4_, 10 MgSO_4_, and 0.5 CaCl_2_ (310 mOsm). Horizontal slices (270 µm) containing RT and TC regions were sectioned using a vibratome (VT1200, Leica) with a slicing speed of 0.07-0.08 mm/s. Slices were incubated on a Brain Slice Keeper 4 (AutoMate Scientific, Berkeley, California) and continuously oxygenated in warm (∼32°C) artificial cerebrospinal fluid (ACSF) containing (in mM): 10 glucose, 26 NaHCO_3_, 2.5 KCl, 1.25 NaHPO_4_, 2 MgSO_4_, 2 CaCl_2_, and 126 NaCl (290 - 300 mOsm) for one hour and then were transferred to room temperature (∼21-23°C) for at least one hour prior to electrophysiological recording.

### Extracellular thalamic oscillations

For oscillation recordings, 400 µm-thick brain slices from Cntnap2^+/+^ and Cntnap2^−/−^ juvenile mice (P21–P25) were placed in a humidified, oxygenated interface recording chamber and perfused with oxygenated artificial cerebrospinal fluid (ACSF) at a flow rate of 2 mL/min, maintained at 32–34°C. To ensure stable perfusion, squares of lens paper were placed beneath and on top of the slice, with a small cutout in the top layer to allow electrode access, and the preparation was secured using platinum bars. To mitigate gradual GABA depletion, L-glutamine (300 µM), a metabolic substrate for GABA synthesis, was added to the ACSF as described by Bryant et al. (2009). For spontaneous oscillation recordings, a tungsten electrode (50–100 kΩ, FHC) was placed in the ventrobasal thalamus (VB), and recordings were conducted continuously for 10 minutes. Optogenetically evoked oscillations were induced by delivering 490 nm blue laser pulses (1 ms duration) to the RT and VB regions every 30 seconds for 10 minutes. For electrical stimulation-induced oscillations, square current pulses were delivered through two parallel tungsten electrodes (50–100 kΩ, FHC) placed 50–100 µm apart in the internal capsule to stimulate both cortical and thalamic axons. Electrical stimuli were 100 µs in duration, 50 V in amplitude, and delivered every 30 seconds for 10 minutes. Extracellular potentials were recorded using a tungsten electrode (50–100 kΩ, FHC) placed in the VB. One experiment was performed per slice, and only slices with a spontaneous bursting frequency of at least five bursts per minute were included in the analyses. Data and statistical analyses were conducted using MATLAB 2023b (MathWorks) and GraphPad Prism 9. Oscillation spike detection and parameter analyses were performed as described by Sohal et al.(*44*). Briefly, spikes were identified as slope deflections greater than three times the threshold, where the threshold was defined as the root-mean-square of background noise during baseline sweeps. Bursts were defined as clusters of spikes (≥3 spikes with <6–10 ms interspike intervals). Oscillations were defined as clusters of bursts (≥2 bursts with <0.6– 1 s interburst intervals). Oscillation duration was measured from the onset of stimulation to the final burst.

### Patch-clamp electrophysiology

Whole-cell patch-clamp recordings of RT neurons were conducted using glass patch pipettes (3–5 MΩ) filled with an intracellular solution containing (in mM): 126 K-gluconate, 4 KCl, 2 Mg-ATP, 0.3 GTP-Tris, 10 phosphocreatine, with pH adjusted to 7.4 using KOH (290 mOsm). For current-clamp recordings, cells were held at a resting membrane potential of approximately –70 mV, with steady intracellular current applied throughout the recording. Experiments were performed in the presence of kynurenic acid and Gabazine (SR-95531) to block fast excitatory and inhibitory synaptic transmission, respectively. Pipette capacitance was compensated, and the bridge balance was adjusted. Evoked action potentials (APs) were elicited by injecting 1-s-duration current steps of increasing amplitude (0–300 pA). The exported traces of APs were further analyzed, and the first and second derivatives (dV/dt and d²V/dt², respectively) were calculated using pClampFit 10.7 and custom MATLAB (MathWorks) code. For voltage-clamp recordings of inhibitory synaptic events, including optogenetically evoked IPSCs (oIPSCs), the pipette solution contained (in mM): 135 CsCl, 10 HEPES, 10 EGTA, 2 MgCl₂, 5 QX-314, with pH adjusted to 7.4 using CsOH (290 mOsm). The membrane potential was held at –70 mV, and signals were recorded using a Multiclamp 700A amplifier with pClamp 10.7 software (Axon Instruments, San Jose, CA) at a sampling rate of 50 kHz, with low-pass filtering at 10 kHz. For measurement of T-type calcium current, pipette capacitance was appropriately compensated. oIPSCs were detected using custom software (Wdetecta, JRH). To block oIPSCs, bath application of 20 µM Gabazine was applied at a holding potential of –70 mV, and no detectable oIPSCs were observed at –5 mV, the expected reversal potential for Cl⁻. Data from unstable recordings were excluded if (1) the series resistance increased by more than 20% from the initial value, or (2) the holding current changed by more than 100 pA.

### Optogenetic stimulation

Mice that underwent stereotaxic injection of AAV5.EF1a.DIO.hChR2(H134R)- eYFP.WPRE.hGH virus into the RT region (AP: − 1.1 to −1.2 mm, ML: ± 2.0 mm, DV: - 3.0 mm) of Cntnap2^+/+^ mice x PV-Cre and Cntnap2^−/−^ mice x PV-Cre mice, were euthanized, and fresh brain slices were prepared for patch-clamp physiology as described above. RT neurons expressing hChR2-eYFP were activated with 475 nm light at an intensity of 1.9 mW/cm². We applied 1 ms pulses of blue LED light at frequencies of 5, 10, 20, 30, and 50 Hz for 5 seconds to stimulate presynaptic release, as previously described.

### In Vivo Fiber photometry

Fiber photometry was used to record fluorescence signals in vivo. For fiber photometry recordings of calcium activity, AAV2/9-EF1a-DIO-GCaMP6s was injected at the following coordinates: AP: −1.10 mm, ML: ±2.00 mm, DV: 3.15 mm, into Cntnap2^+/+^ x PV-cre and Cntnap2^−/−^ x PV-cre mice. Optic fiber cannulae with a 200 µm core, 0.39 NA, and 1.25 mm diameter ferrule (RWD Life Science) were implanted at AP: −1.10 mm, ML: ±2.00 mm, and DV: 3.00 mm from bregma for the RT. The 4.0 mm long optic fiber cannulae were used for RT and fixed to the skull with dental cement (C&B Metabond, Parkell). Animals were allowed to recover from surgery for 3 weeks before the recording experiments. In vivo fiber photometry recordings were performed using a dual-color multichannel fiber photometry system (R811, RWD Life Science) with a low-autofluorescence 1–4 fan-out bundled fiber patch cord with a 200 µm core, 0.39 NA, and 1.25 mm diameter ferrule. GCaMP6s excitation was at 470 nm, and 410 nm was used for its isosbestic control. Green fluorescence emission was recorded at 30 Hz. The data from fiber photometry were processed and analyzed as follows: the photometry signal value, *F*, was calculated as *F_470_/F_410_*. *ΔF/F* was then derived using the formula *ΔF/F = (F - F₀) / F₀,* where *F₀* is the median of the photometry signal. Only calcium signals greater than 3 standard deviations (>3 SD) were treated as events, detected using the findpeaks function in MATLAB.

### Quantification and Statistical analysis

All bar graphs indicate the mean and all error bars represent ± standard error of the mean (SEM). Statistical analyses were performed using GraphPad Prism 9 (GraphPad Software, La Jolla, CA), Excel (Microsoft), and Custom MATLAB programs (MathWorks, Natick, MA). Normally distributed data with equal variance were analyzed using t-tests (two-tailed, unpaired or paired), one-way ANOVA, or two-way ANOVA, followed by Tukey’s multiple comparison test. Statistical values are denoted as follows: *p < 0.05, **p < 0.01, and ***p < 0.001.

## Acknowledgments

We thank all current lab members for their feedback during the entire project. We also thank Nicole Agranonik for managing mouse lines, laboratory equipment and supplies.

## Author contributions

Conceptualization: S.S.J and J.R.H. Methodology: S.S.J and J.R.H. Investigation: S.S.J, F.T and J.R.H. Visualization: S.S.J. Supervision: J.R.H. Writing—original draft: S.S.J. Writing—review & editing: S.S.J and J.R.H. Funding acquisition: J.R.H.

## Data and materials availability

All data needed to evaluate the conclusions in the paper are present in the paper and/or the Supplementary Materials.

---

**Fig. S1.**
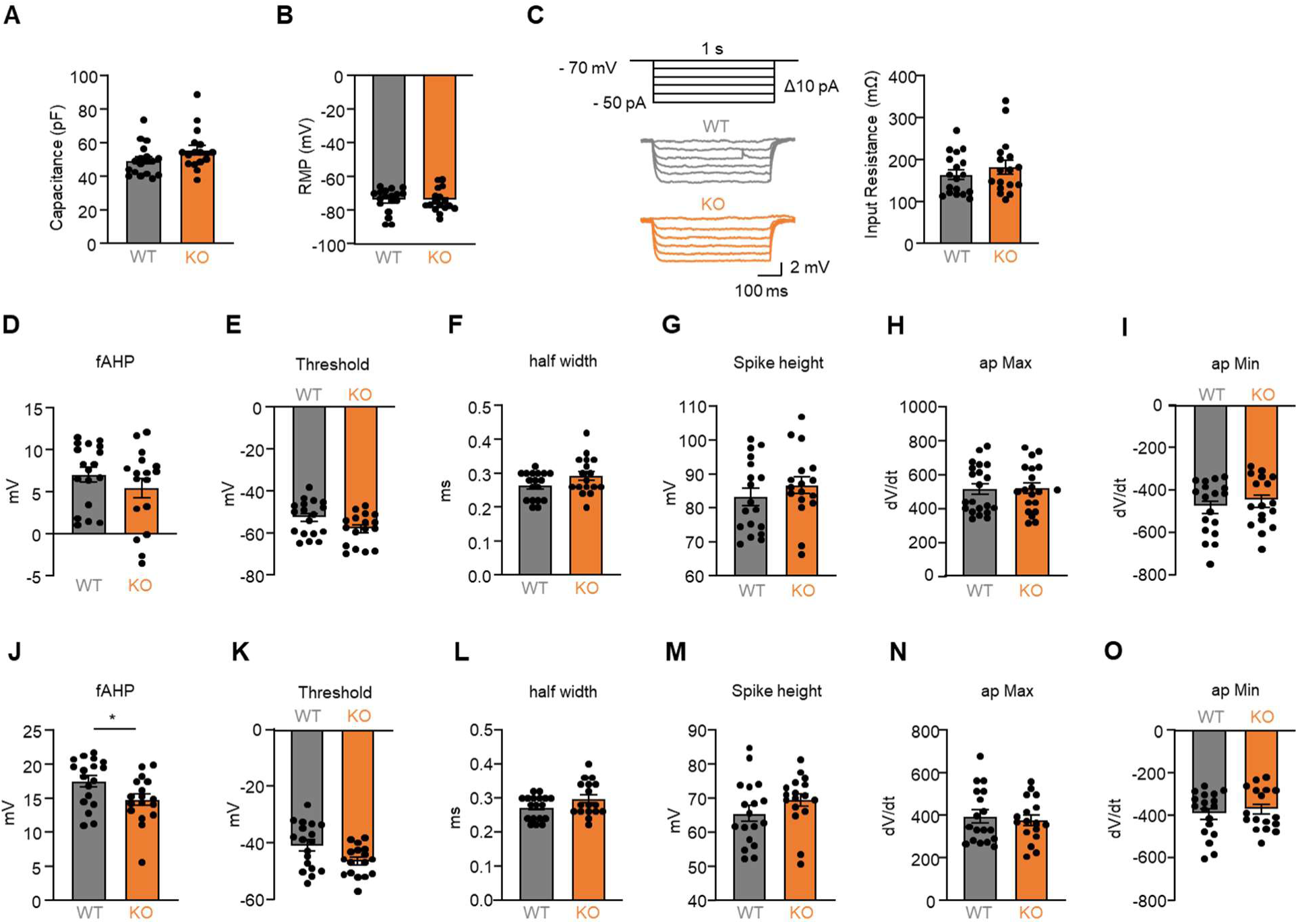
Passive properties and AP properties in RT of Cntnap2^+/+^ and Cntnap2^−/−^ mice. **(A-B)** Quantification of membrane capacitance and resting membrane potential (RMP) in RT of Cntnap2^+/+^ and Cntnap2^−/−^ mice. **(C)** Protocol and representative traces (left) showing voltage deflections elicited by – 50 pA to –10 pA intracellular current injections and (right) quantification of input resistance (mΩ) from RT of Cntnap2^+/+^ and Cntnap2^−/−^ mice. **(D-I)** Quantification of AP properties, including fAHP, AP threshold, half width, spike height, max dV/dt, and min dV/dt from 1^st^ rheobase spike from RT of Cntnap2^+/+^ and Cntnap2^−/−^ mice. **(J-O)** As with (D-I), but for last spike from RT of Cntnap2^+/+^ and Cntnap2^−/−^ mice. **(A-O)** n = 18 cells from 4 mice for Cntnap2^+/+^ and n =17 cells from 4 mice for Cntnap2^−/−^. The statistical tests involved an unpaired two-tailed t-test (A-O). Data are represented as mean values ± SEM. *P*-values in figure panels: *p < 0.05.

**Fig. S2.**
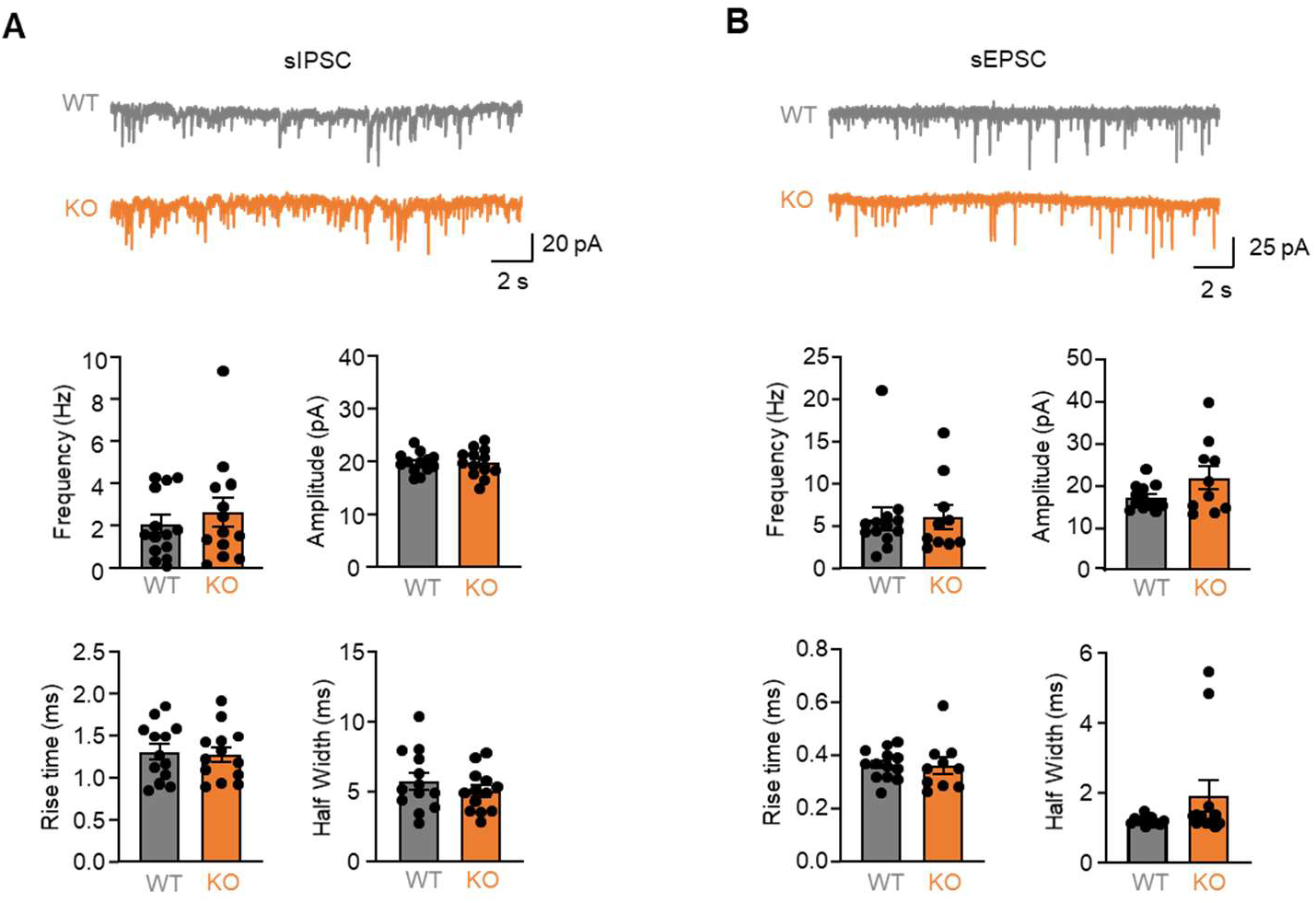
sIPSCs and sEPSCs in RT of Cntnap2^+/+^ and Cntnap2^−/−^ mice. **(A)** Representative traces showing spontaneous IPSCs (top) and quantification of sIPSC frequency (Hz), amplitude (pA), rise time (ms), and half-width (ms) (bottom) from RT of Cntnap2^+/+^ and Cntnap2^−/−^ mice. (n = 13 cells from 3 mice for Cntnap2^+/+^ and n = 13 cells from 3 mice for Cntnap2^−/−^). **(B)** Representative traces showing spontaneous EPSCs (top) and quantification of sEPSC frequency (Hz), amplitude (pA), rise time (ms), and half-width (ms) (bottom) from RT of Cntnap2^+/+^ and Cntnap2^−/−^ mice. (n =13 cells from 3 mice for Cntnap2^+/+^ and n =10 cells from 3 mice for Cntnap2^−/−^). The statistical tests involved an unpaired two-tailed t-test (A,B). Data are represented as mean values ± SEM.

**Fig. S3.**
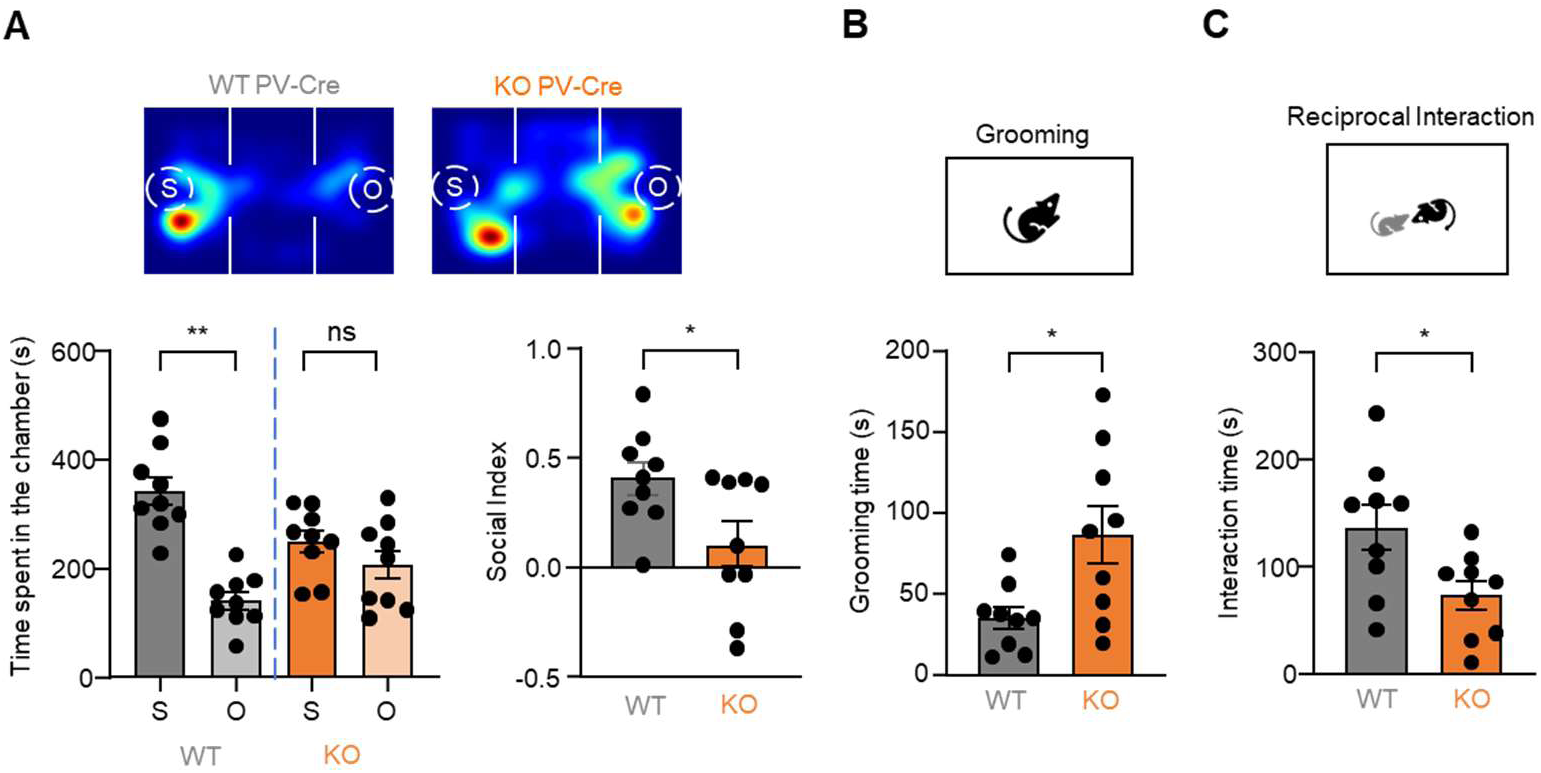
Confirmation of deficits in ASD-related behaviors in Cntnap2^+/+^ and Cntnap2^−/−^ mice PV-Cre mice. **(A)** Representative heatmaps showing location in social preference tests (top), and time spent in the chamber (bottom, left) and social index (bottom, right) in Cntnap2^+/+^ and Cntnap2^−/−^ PV-Cre mice. **(B)** Quantification of grooming time in Cntnap2^+/+^ and Cntnap2^−/−^ PV-cre mice. **(C)** Quantification of interaction time in Cntnap2^+/+^ and Cntnap2^−/−^ PV-cre mice. **(A-C)** n = 9 mice for Cntnap2^+/+^ PV-cre and n =9 mice for Cntnap2^−/−^ PV-cre. The statistical tests involved an unpaired two-tailed t-test (A-C). Data are represented as mean values ± SEM. *P*-values in figure panels: *p < 0.05, **p < 0.01, ns, not significant.

**Fig. S4.**
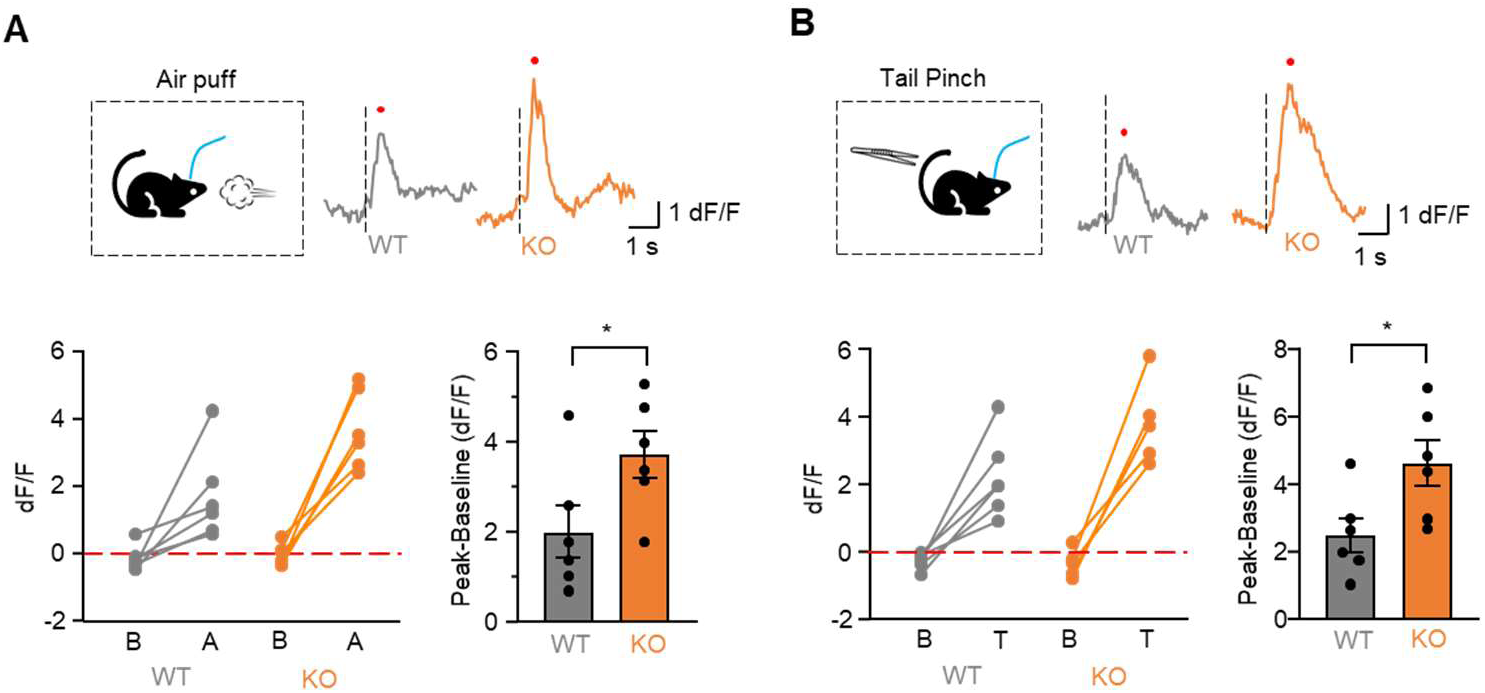
Ca^2+^ response to acute aversive stimuli in Cntnap2^+/+^ and Cntnap2^−/−^ mice. **(A)** Schematic illustrating the air puff (top, left), representative Ca^2+^ signal (top, right), and quantification of peak dF/F and change in dF/F (bottom) during baseline and response to air puff in Cntnap2^+/+^ and Cntnap2^−/−^ mice. **(B)** Schematic illustrating the tail pinch (top left), representative Ca^2+^ signal (top, right), and quantification of peak dF/F and change of dF/F (bottom) during baseline and response to tail pinch in Cntnap2^+/+^ and Cntnap2^−/−^ mice. **(A-B)** n = 6 mice for Cntnap2^+/+^ and n = 6 mice for Cntnap2^−/−^. The statistical tests involved a paired two-tailed t-test (A-B). Data are represented as mean values ± SEM. *P*-values in figure panels: *p < 0.05.

**Fig. S5.**
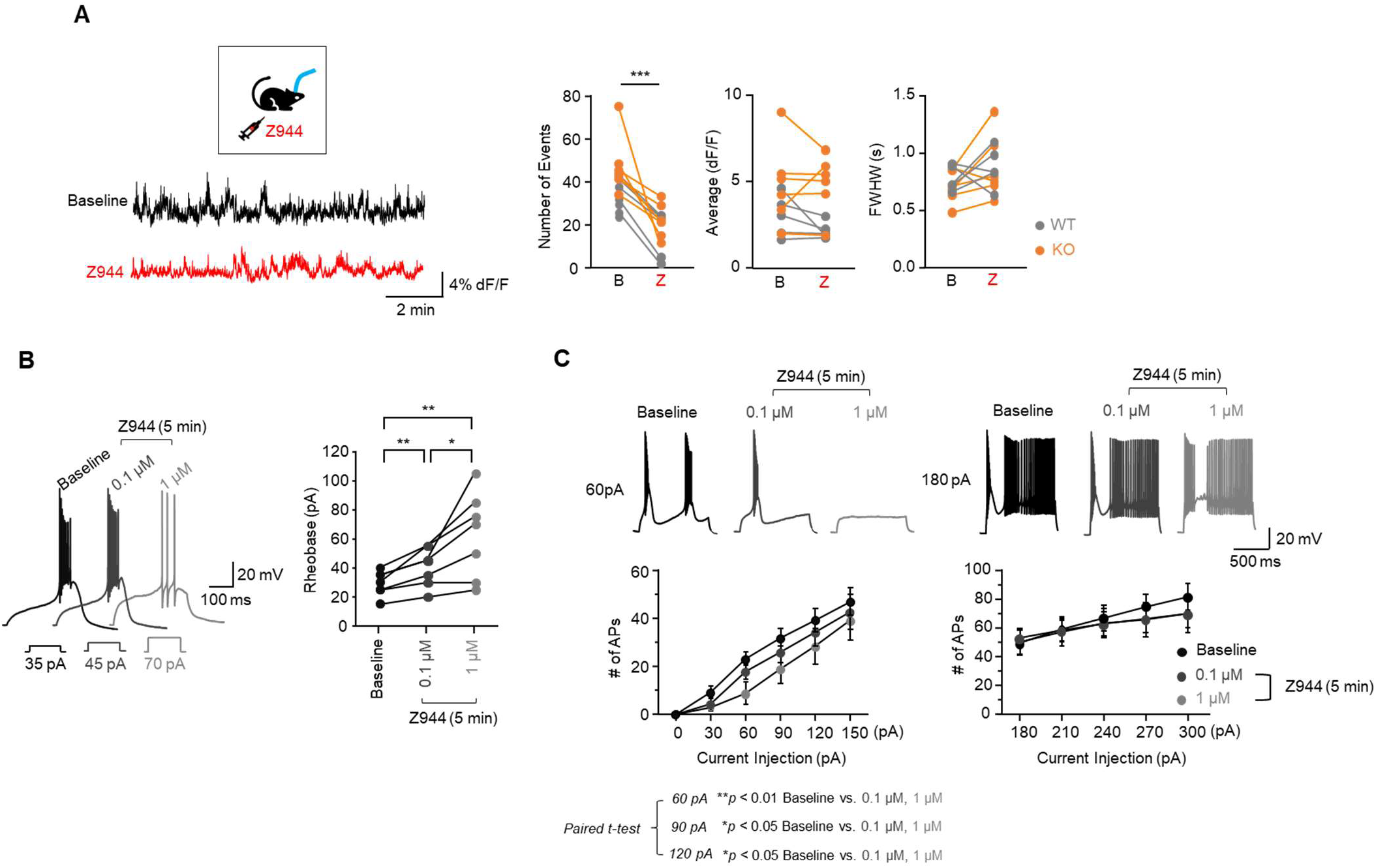
Z944 inhibits in vivo RT population activity, elevates rheobase, and reduces excitability in RT. **(A)** Schematic (left, top) illustrating experiment, representative fiber photometry signals (left, bottom) of spontaneous Ca^2+^ fluctuations during baseline and Z944 condition, (right) quantification of number of events, average (dF/F), and FWHM in Ca^2+^ dynamics during baseline and Z944 condition (right). (n = 5 mice for Cntnap2^+/+^ and n = 6 mice for Cntnap2^−/−^). **(B)** Representative traces (left) showing burst APs elicited by rheobase currents and graphs (right) showing quantification of rheobase currents (pA) during baseline, 0.1 µM, and 1 µM Z944 for 5 min incubation. **(C)** Representative traces (top) showing burst APs elicited by 60 pA (top, left) and 180 pA (top, right) injection and graph (bottom) and quantification of the number of evoked APs with weak (bottom, left) and strong current injection (bottom, right) during baseline, 0.1 µM, and 1 µM Z944 for 5 min incubation. The statistical tests involved a paired two-tailed t-test. Data are represented as mean values ± SEM. *P*-values in figure panels: *p < 0.05. ***p < 0.001.

**Fig. S6.**
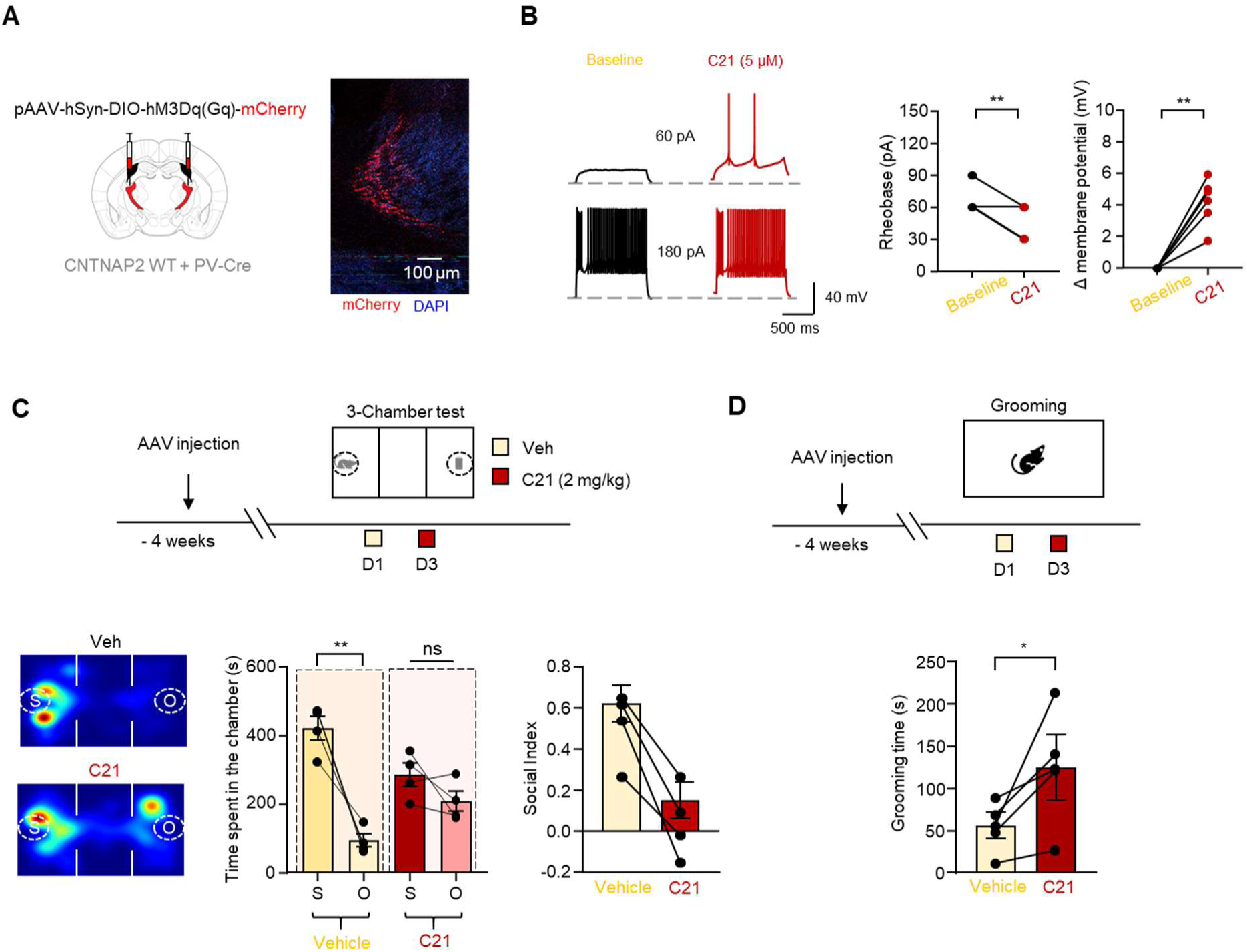
Chemogenetic activation of RT impairs social preference and grooming behavior in Cntnap2^+/+^ mice. **(A)** Schematic describing stereotaxic injection of pAAV-hSyn-DIO-hM3D(Gq)-mCherry into RT (left) and red fluorescence representing expression of mCherry (right). **(B)** Representative traces showing APs (left) elicited by rheobase currents and quantification of c21 induced changes in membrane potential and rheobase current (right). (n = 6 cells from 2 mice). **(C)** Experimental strategy for evaluating the effect of chemogenetic RT activation on social preference (top) and representative heatmaps showing location during the social preference tests (bottom, left) and time spent in the chamber (bottom, middle) and the social index. (bottom, right) in Cntnap2^+/+^ and Cntnap2^−/−^mice treated with vehicle or C21. **(D)** Experimental strategy for evaluating the effect of chemogenetic RT activation on grooming behavior (top) and quantification of grooming time (bottom) in Cntnap2^+/+^ and Cntnap2^−/−^ mice treated with vehicle or C21. The statistical tests involved a paired two-tailed t-test (B, D, F). Data are represented as mean values ± SEM. *P*-values in figure panels: *p < 0.05, **p < 0. 01, ***p < 0.001, ns, not significant.

**Fig. S7.**
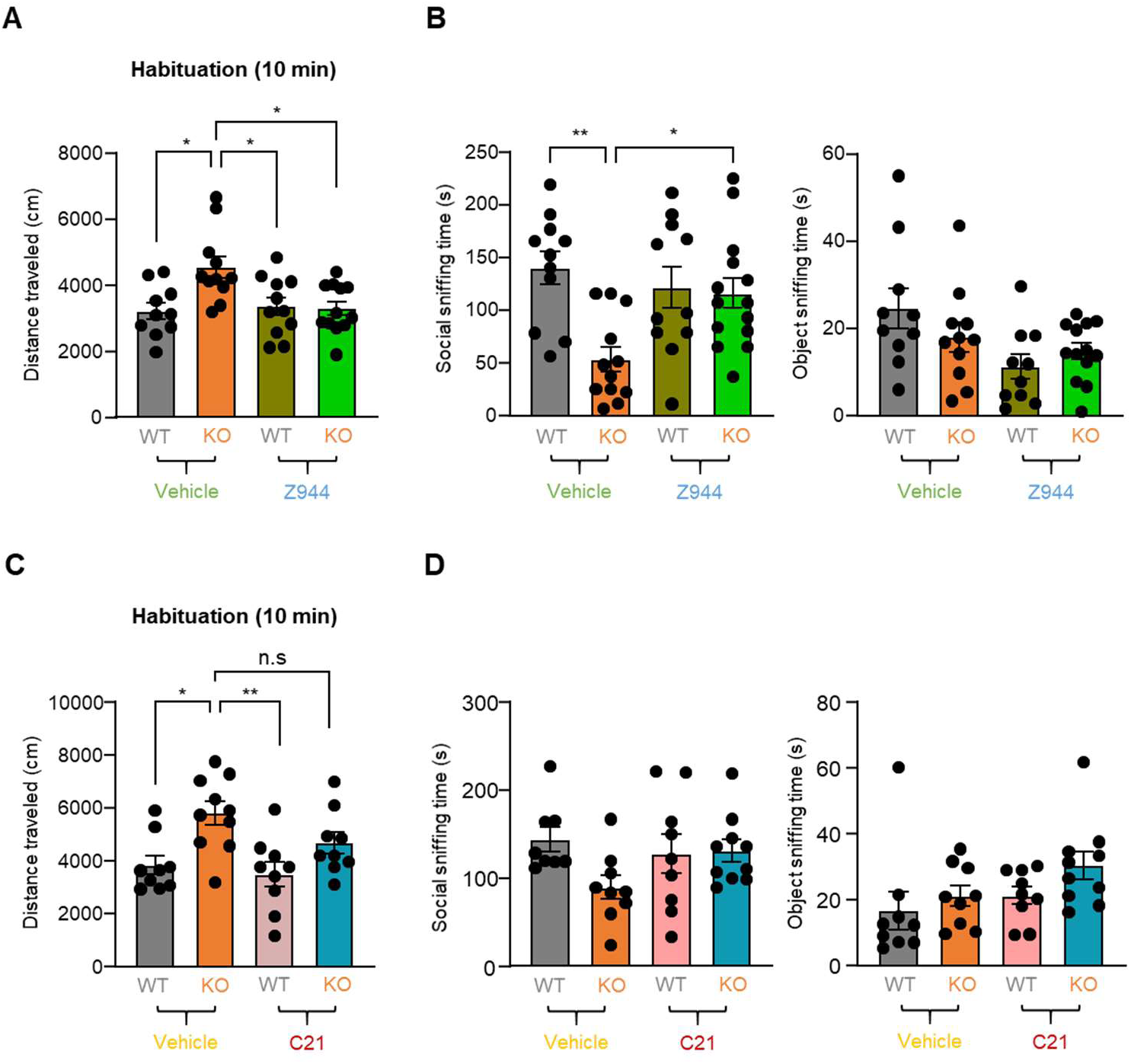
Locomotor activity during habituation period and sniffing time (social and object) following inhibition of either T-type calcium channels or activation of hM4D (Gi) in RT neurons. **(A)** Quantification of total distance traveled during habituation period (10 min) in Cntnap2^+/+^ and Cntnap2^−/−^mice treated with vehicle or Z944. (n = 10 mice for Cntnap2^+/+^- Veh, n = 11 for Cntnap2^−/−^-Veh, n = 11 for Cntnap2^+/+^ -Z944, n = 12 for Cntnap2^−/−^-Z944). **(B)** Quantification of social (left) and object (right) sniffing time over 10 min in Cntnap2^+/+^ and Cntnap2^−/−^mice treated with vehicle or Z944. (n = 11 mice for Cntnap2^+/+^-Veh, n = 11 for Cntnap2^−/−^-Veh, n = 10 for Cntnap2^+/+^-Z944, n = 13 for Cntnap2^−/−^-Z944). **(C)** Quantification of total distance traveled during habituation period (10 min) in Cntnap2^+/+^ and Cntnap2^−/−^ mice treated with vehicle or C21. (n = 9 mice for Cntnap2^+/+^-Veh, n = 10 for Cntnap2^−/−^-Veh, n = 9 for Cntnap2^+/+^-C21, n = 9 for Cntnap2^−/−^-C21). **(D)** Quantification of social (left) and object (right) sniffing time over 10 min in Cntnap2^+/+^ and Cntnap2^−/−^mice treated with vehicle or C21. (n = 8 mice for Cntnap2^+/+^-Veh, n = 9 for Cntnap2^−/−^-Veh, n = 9 for Cntnap2^+/+^-C21, n = 10 for Cntnap2^−/−^-C21). The statistical tests involved two-way ANOVA with Tukey’s test (A-B). Data are represented as mean values ± SEM. *P*-values in figure panels: *p < 0.05, **p < 0.01, n.s, not significant.

## References

1. O. Peñagarikano, B. S. Abrahams, E. I. Herman, K. D. Winden, A. Gdalyahu, H. Dong, L. I. Sonnenblick, R. Gruver, J. Almajano, A. Bragin, P. Golshani, J. T. Trachtenberg, E. Peles, D. H. Geschwind, Absence of CNTNAP2 Leads to Epilepsy, Neuronal Migration Abnormalities, and Core Autism-Related Deficits. Cell 147, 235–246 (2011).

2. K. R. Cording, E. M. Tu, H. Wang, A. H. Agopyan-Miu, H. S. Bateup, Cntnap2 loss drives striatal neuron hyperexcitability and behavioral inflexibility. [Preprint] (2024). 10.7554/eLife.100162.1.

3. W. E. Jang, J. H. Park, G. Park, G. Bang, C. H. Na, J. Y. Kim, K.-Y. Kim, K. P. Kim, C. Y. Shin, J.-Y. An, Y.-S. Lee, M.-S. Kim, Cntnap2-dependent molecular networks in autism spectrum disorder revealed through an integrative multi-omics analysis. Mol. Psychiatry 28, 810–821 (2023).

4. H. C. Wang, D. E. Feldman, Degraded tactile coding in the Cntnap2 mouse model of autism. Cell Rep. 43, 114612 (2024).

5. E. Lauber, F. Filice, B. Schwaller, Dysregulation of Parvalbumin Expression in the Cntnap2−/− Mouse Model of Autism Spectrum Disorder. Front. Mol. Neurosci. 11, 262 (2018).

6. M. J. Gandal, J. R. Haney, B. Wamsley, C. X. Yap, S. Parhami, P. S. Emani, N. Chang, G. T. Chen, G. D. Hoftman, D. De Alba, G. Ramaswami, C. L. Hartl, A. Bhattacharya, C. Luo, T. Jin, D. Wang, R. Kawaguchi, D. Quintero, J. Ou, Y. E. Wu, N. N. Parikshak, V. Swarup, T. G. Belgard, M. Gerstein, B. Pasaniuc, D. H. Geschwind, Broad transcriptomic dysregulation occurs across the cerebral cortex in ASD. Nature 611, 532– 539 (2022).

7. H. Won, H.-R. Lee, H. Y. Gee, W. Mah, J.-I. Kim, J. Lee, S. Ha, C. Chung, E. S. Jung, Y. S. Cho, S.-G. Park, J.-S. Lee, K. Lee, D. Kim, Y. C. Bae, B.-K. Kaang, M. G. Lee, E. Kim, Autistic-like social behaviour in Shank2-mutant mice improved by restoring NMDA receptor function. Nature 486, 261–265 (2012).

8. M. T. Lazaro, J. Taxidis, T. Shuman, I. Bachmutsky, T. Ikrar, R. Santos, G. M. Marcello, A. Mylavarapu, S. Chandra, A. Foreman, R. Goli, D. Tran, N. Sharma, M. Azhdam, H. Dong, K. Y. Choe, O. Peñagarikano, S. C. Masmanidis, B. Rácz, X. Xu, D. H. Geschwind, P. Golshani, Reduced Prefrontal Synaptic Connectivity and Disturbed Oscillatory Population Dynamics in the CNTNAP2 Model of Autism. Cell Rep. 27, 2567–2578.e6 (2019).

9. R. Paterno, J. R. Marafiga, H. Ramsay, T. Li, K. A. Salvati, S. C. Baraban, Hippocampal gamma and sharp-wave ripple oscillations are altered in a Cntnap2 mouse model of autism spectrum disorder. Cell Rep. 37, 109970 (2021).

10. J. K. Kern, M. H. Trivedi, C. R. Garver, B. D. Grannemann, A. A. Andrews, J. S. Savla, D. G. Johnson, J. A. Mehta, J. L. Schroeder, The pattern of sensory processing abnormalities in autism. Autism 10, 480–494 (2006).

11. E. J. Marco, L. B. N. Hinkley, S. S. Hill, S. S. Nagarajan, Sensory Processing in Autism: A Review of Neurophysiologic Findings: *Pediatr*. Res. 69, 48R–54R (2011).

12. M. O. Mazurek, K. Sohl, Sleep and Behavioral Problems in Children with Autism Spectrum Disorder. J. Autism Dev. Disord. 46, 1906–1915 (2016).

13. M. C. Souders, T. B. A. Mason, O. Valladares, M. Bucan, S. E. Levy, D. S. Mandell, T. E. Weaver, J. Pinto-Martin, Sleep Behaviors and Sleep Quality in Children with Autism Spectrum Disorders. 32 (2009).

14. S. Lukmanji, S. A. Manji, S. Kadhim, K. M. Sauro, E. C. Wirrell, C.-S. Kwon, N. Jetté, The co-occurrence of epilepsy and autism: A systematic review. Epilepsy Behav. 98, 238–248 (2019).

15. J. B. Ewen, A. R. Marvin, K. Law, P. H. Lipkin, Epilepsy and Autism Severity: A Study of 6,975 Children. Autism Res. 12, 1251–1259 (2019).

16. A. Nair, J. M. Treiber, D. K. Shukla, P. Shih, R.-A. Müller, Impaired thalamocortical connectivity in autism spectrum disorder: a study of functional and anatomical connectivity. Brain 136, 1942–1955 (2013).

17. M. Schuetze, M. T. M. Park, I. Y. Cho, F. P. MacMaster, M. M. Chakravarty, S. L. Bray, Morphological Alterations in the Thalamus, Striatum, and Pallidum in Autism Spectrum Disorder. Neuropsychopharmacology 41, 2627–2637 (2016).

18. T. Iidaka, T. Kogata, Y. Mano, H. Komeda, Thalamocortical Hyperconnectivity and Amygdala-Cortical Hypoconnectivity in Male Patients With Autism Spectrum Disorder. Front. Psychiatry 10, 252 (2019).

19. B. Guo, T. Liu, S. Choi, H. Mao, W. Wang, K. Xi, C. Jones, N. D. Hartley, D. Feng, Q. Chen, Y. Liu, R. D. Wimmer, Y. Xie, N. Zhao, J. Ou, M. A. Arias-Garcia, D. Malhotra, Y. Liu, S. Lee, S. Pasqualoni, R. J. Kast, M. Fleishman, M. M. Halassa, S. Wu, Z. Fu, Restoring thalamocortical circuit dysfunction by correcting HCN channelopathy in Shank3 mutant mice. Cell Rep. Med. 5, 101534 (2024).

20. A. Clemente-Perez, S. R. Makinson, B. Higashikubo, S. Brovarney, F. S. Cho, A. Urry, S. S. Holden, M. Wimer, C. Dávid, L. E. Fenno, L. Acsády, K. Deisseroth, J. T. Paz, Distinct Thalamic Reticular Cell Types Differentially Modulate Normal and Pathological Cortical Rhythms. Cell Rep. 19, 2130–2142 (2017).

21. R. I. Martinez-Garcia, B. Voelcker, J. B. Zaltsman, S. L. Patrick, T. R. Stevens, B. W. Connors, S. J. Cruikshank, Two dynamically distinct circuits drive inhibition in the sensory thalamus. Nature 583, 813–818 (2020).

22. Y. Li, V. G. Lopez-Huerta, X. Adiconis, K. Levandowski, S. Choi, S. K. Simmons, M. A. Arias-Garcia, B. Guo, A. Y. Yao, T. R. Blosser, R. D. Wimmer, T. Aida, A. Atamian, T. Naik, X. Sun, D. Bi, D. Malhotra, C. C. Hession, R. Shema, M. Gomes, T. Li, E. Hwang, A. Krol, M. Kowalczyk, J. Peça, G. Pan, M. M. Halassa, J. Z. Levin, Z. Fu, G. Feng, Distinct subnetworks of the thalamic reticular nucleus. Nature 583, 819–824 (2020).

23. S. Ahrens, S. Jaramillo, K. Yu, S. Ghosh, G.-R. Hwang, R. Paik, C. Lai, M. He, Z. J. Huang, B. Li, ErbB4 regulation of a thalamic reticular nucleus circuit for sensory selection. Nat. Neurosci. 18, 104–111 (2015).

24. G. Vantomme, Z. Rovó, R. Cardis, E. Béard, G. Katsioudi, A. Guadagno, V. Perrenoud, L. M. J. Fernandez, A. Lüthi, A Thalamic Reticular Circuit for Head Direction Cell Tuning and Spatial Navigation. Cell Rep. 31, 107747 (2020).

25. J.-H. Lee, C.-F. V. Latchoumane, J. Park, J. Kim, J. Jeong, K.-H. Lee, H.-S. Shin, The rostroventral part of the thalamic reticular nucleus modulates fear extinction. Nat. Commun. 10, 4637 (2019).

26. L. D. Lewis, J. Voigts, F. J. Flores, L. I. Schmitt, M. A. Wilson, M. M. Halassa, E. N. Brown, Thalamic reticular nucleus induces fast and local modulation of arousal state. eLife 4, e08760 (2015).

27. M. F. Wells, Thalamic reticular impairment underlies attention deficit in Ptchd1Y/- mice.

28. X. Zhang, X. Yu, M. Tuo, Z. Zhao, J. Wang, T. Jiang, M. Zhang, Y. Wang, Y. Sun, Parvalbumin neurons in the anterior nucleus of thalamus control absence seizures. Epilepsia Open 8, 1002–1012 (2023).

29. P.-F. Liu, Y. Wang, L. Xu, A.-F. Xiang, M.-Z. Liu, Y.-B. Zhu, X. Jia, R. Zhang, J.-B. Li, L. Zhang, D. Mu, Modulation of itch and pain signals processing in ventrobasal thalamus by thalamic reticular nucleus. iScience 25, 103625 (2022).

30. J. Huguenard, D. Prince, A novel T-type current underlies prolonged Ca(2+)- dependent burst firing in GABAergic neurons of rat thalamic reticular nucleus. J. Neurosci. 12, 3804–3817 (1992).

31. C. L. Cox, J. R. Huguenard, D. A. Prince, Nucleus reticularis neurons mediate diverse inhibitory effects in thalamus. Proc. Natl. Acad. Sci. 94, 8854–8859 (1997).

32. C. L. Cox, J. R. Huguenard, D. A. Prince, Heterogeneous axonal arborizations of rat thalamic reticular neurons in the ventrobasal nucleus. J. Comp. Neurol. 366, 416–430 (1996).

33. V. S. Sohal, M. M. Huntsman, J. R. Huguenard, Reciprocal Inhibitory Connections Regulate the Spatiotemporal Properties of Intrathalamic Oscillations. J. Neurosci. 20, 1735–1745 (2000).

34. M. M. Halassa, J. H. Siegle, J. T. Ritt, J. T. Ting, G. Feng, C. I. Moore, Selective optical drive of thalamic reticular nucleus generates thalamic bursts and cortical spindles. Nat. Neurosci. 14, 1118–1120 (2011).

35. C. M. Schofield, M. Kleiman-Weiner, U. Rudolph, J. R. Huguenard, A gain in GABA_A_ receptor synaptic strength in thalamus reduces oscillatory activity and absence seizures. Proc. Natl. Acad. Sci. 106, 7630–7635 (2009).

36. P. M. Fogerson, J. R. Huguenard, Tapping the Brakes: Cellular and Synaptic Mechanisms that Regulate Thalamic Oscillations. Neuron 92, 687–704 (2016).

37. C. El Khoueiry, J.-H. Cabungcal, Z. Rovó, M. Fournier, K. Q. Do, P. Steullet, Developmental oxidative stress leads to T-type Ca2+ channel hypofunction in thalamic reticular nucleus of mouse models pertinent to schizophrenia. Mol. Psychiatry 27, 2042– 2051 (2022).

38. X. Zhu, J.-H. Cabungcal, M. Cuenod, D. L. Uliana, K. Q. Do, A. A. Grace, Thalamic reticular nucleus impairments and abnormal prefrontal control of dopamine system in a developmental model of schizophrenia: prevention by N-acetylcysteine. Mol. Psychiatry 26, 7679–7689 (2021).

39. X.-Y. Wang, X. Xu, R. Chen, W.-B. Jia, P.-F. Xu, X.-Q. Liu, Y. Zhang, X.-F. Liu, Y. Zhang, The thalamic reticular nucleus-lateral habenula circuit regulates depressive-like behaviors in chronic stress and chronic pain. Cell Rep. 42, 113170 (2023).

40. A. M. Thomas, M. D. Schwartz, M. D. Saxe, T. S. Kilduff, Cntnap2 Knockout Rats and Mice Exhibit Epileptiform Activity and Abnormal Sleep–Wake Physiology. Sleep 40 (2017).

41. A. N. Mohapatra, R. Jabarin, N. Ray, S. Netser, S. Wagner, Impaired emotion recognition in Cntnap2-deficient mice is associated with hyper-synchronous prefrontal cortex neuronal activity. Mol. Psychiatry, 1–13 (2024).

42. S. Jurgensen, P. E. Castillo, Selective Dysregulation of Hippocampal Inhibition in the Mouse Lacking Autism Candidate Gene *CNTNAP2*. J. Neurosci. 35, 14681–14687 (2015).

43. M. S. Dawson, K. Gordon-Fleet, L. Yan, V. Tardos, H. He, K. Mui, S. Nawani, Z. Asgarian, M. Catani, C. Fernandes, U. Drescher, Sexual dimorphism in the social behaviour of Cntnap2-null mice correlates with disrupted synaptic connectivity and increased microglial activity in the anterior cingulate cortex. *Commun*. Biol. 6, 846 (2023).

44. V. S. Sohal, J. R. Huguenard, Inhibitory Interconnections Control Burst Pattern and Emergent Network Synchrony in Reticular Thalamus. J. Neurosci. 23, 8978–8988 (2003).

45. S. S. Jeste, R. Tuchman, Autism Spectrum Disorder and Epilepsy: Two Sides of the Same Coin? J. Child Neurol. 30, 1963–1971 (2015).

46. S. N. Mandhane, K. Aavula, T. Rajamannar, Timed pentylenetetrazol infusion test: A comparative analysis with s.c.PTZ and MES models of anticonvulsant screening in mice. Seizure - Eur. J. Epilepsy 16, 636–644 (2007).

47. E. L. Hill-Yardin, A. Argyropoulos, S. Hosie, G. Rind, P. Anderson, A. J. Hannan, T. J. O’Brien, Reduced susceptibility to induced seizures in the Neuroligin-3R451C mouse model of autism. Neurosci. Lett. 589, 57–61 (2015).

48. G. M. Jacobson, L. J. Voss, S. M. Melin, J. P. Mason, R. T. Cursons, D. A. Steyn-Ross, M. L. Steyn-Ross, J. W. Sleigh, Connexin36 knockout mice display increased sensitivity to pentylenetetrazol-induced seizure-like behaviors. Brain Res. 1360, 198–204 (2010).

49. A. Ruszczak, P. Poznański, A. Leśniak, M. Łazarczyk, D. Skiba, A. Nawrocka, K. Gaweł, J. Paszkiewicz, M.-E. Mickael, M. Sacharczuk, Susceptibility to Pentylenetetrazole-Induced Seizures in Mice with Distinct Activity of the Endogenous Opioid System. Int. J. Mol. Sci. 25, 6978 (2024).

50. J. V. Erum, D. V. Dam, P. P. D. Deyn, PTZ-induced seizures in mice require a revised Racine scale. Epilepsy Behav. 95, 51–55 (2019).

51. S. Ritter-Makinson, A. Clemente-Perez, B. Higashikubo, F. S. Cho, S. S. Holden, E. Bennett, A. Chkhaidze, O. H. J. Eelkman Rooda, M.-C. Cornet, F. E. Hoebeek, K. Yamakawa, M. R. Cilio, B. Delord, J. T. Paz, Augmented Reticular Thalamic Bursting and Seizures in Scn1a-Dravet Syndrome. Cell Rep. 26, 54–64.e6 (2019).

52. C. D. Makinson, B. S. Tanaka, J. M. Sorokin, J. C. Wong, C. A. Christian, A. L. Goldin, A. Escayg, J. R. Huguenard, Regulation of Thalamic and Cortical Network Synchrony by Scn8a. Neuron 93, 1165–1179.e6 (2017).

53. J. T. Paz, A. S. Bryant, K. Peng, L. Fenno, O. Yizhar, W. N. Frankel, K. Deisseroth, J. R. Huguenard, A new mode of corticothalamic transmission revealed in the Gria4−/− model of absence epilepsy. Nat. Neurosci. 14, 1167–1173 (2011).

54. C. L. Kyuyoung, J. R. Huguenard, Modulation of Short-Term Plasticity in the Corticothalamic Circuit by Group III Metabotropic Glutamate Receptors. J. Neurosci. 34, 675–687 (2014).

55. A. C. Lu, C. K. Lee, M. Kleiman-Weiner, B. Truong, M. Wang, J. R. Huguenard, M. P. Beenhakker, Nonlinearities between inhibition and T-type calcium channel activity bidirectionally regulate thalamic oscillations. eLife 9, e59548 (2020).

56. R. B. Jacobsen, D. Ulrich, J. R. Huguenard, GABAB and NMDA Receptors Contribute to Spindle-Like Oscillations in Rat Thalamus In Vitro. J. Neurophysiol. 86, 1365–1375 (2001).

57. G. Hou, A. G. Smith, Z.-W. Zhang, Lack of Intrinsic GABAergic Connections in the Thalamic Reticular Nucleus of the Mouse. J. Neurosci. 36, 7246–7252 (2016).

58. M. Abdelaal, M. Midorikawa, T. Suzuki, K. Kobayashi, N. Takata, M. Miyata, M. Mimura, K. Tanaka, Dysfunction of parvalbumin-expressing cells in the thalamic reticular nucleus induces cortical spike-and-wave discharges and an unconscious state. Brain Commun. 4 (2022).

59. M. Wöhr, D. Orduz, P. Gregory, H. Moreno, U. Khan, K. J. Vörckel, D. P. Wolfer, H. Welzl, D. Gall, S. N. Schiffmann, B. Schwaller, Lack of parvalbumin in mice leads to behavioral deficits relevant to all human autism core symptoms and related neural morphofunctional abnormalities. Transl. Psychiatry 5, e525–e525 (2015).

60. F. Filice, K. J. Vörckel, A. Ö. Sungur, M. Wöhr, B. Schwaller, Reduction in parvalbumin expression not loss of the parvalbumin-expressing GABA interneuron subpopulation in genetic parvalbumin and shank mouse models of autism. Mol. Brain 9, 10 (2016).

61. G. Calfa, W. Li, J. M. Rutherford, L. Pozzo-Miller, Excitation/inhibition imbalance and impaired synaptic inhibition in hippocampal area CA3 of *Mecp2* knockout mice: E/I Imbalance In Mecp2 Knockout Hippocampus. Hippocampus 25, 159–168 (2015).

62. A. Caballero, E. Flores-Barrera, D. R. Thomases, K. Y. Tseng, Downregulation of parvalbumin expression in the prefrontal cortex during adolescence causes enduring prefrontal disinhibition in adulthood. Neuropsychopharmacology 45, 1527–1535 (2020).

63. N. D. Hartley, A. Krol, S. Choi, N. Rome, K. Levandowski, S. Pasqualoni, C. Jones, J. Tian, S. Lee, H. Lee, R. Kast, G. Feng, Z. Fu, Distinct structural and functional connectivity of genetically segregated thalamoreticular subnetworks. Cell Rep. 43, 115037 (2024).

64. W. Ding, L. Yang, E. Shi, B. Kim, S. Low, K. Hu, L. Gao, P. Chen, W. Ding, D. Borsook, A. Luo, J. H. Choi, C. Wang, O. Akeju, J. Yang, C. Ran, K. L. Schreiber, J. Mao, Q. Chen, G. Feng, S. Shen, The endocannabinoid N-arachidonoyl dopamine is critical for hyperalgesia induced by chronic sleep disruption. Nat. Commun. 14, 6696 (2023).

65. E. Tringham, K. L. Powell, S. M. Cain, K. Kuplast, J. Mezeyova, M. Weerapura, C. Eduljee, X. Jiang, P. Smith, J.-L. Morrison, N. C. Jones, E. Braine, G. Rind, M. Fee-Maki, D. Parker, H. Pajouhesh, M. Parmar, T. J. O’Brien, T. P. Snutch, T-Type Calcium Channel Blockers That Attenuate Thalamic Burst Firing and Suppress Absence Seizures. Sci. Transl. Med. 4 (2012).

66. C. Cascio, F. McGlone, S. Folger, V. Tannan, G. Baranek, K. A. Pelphrey, G. Essick, Tactile Perception in Adults with Autism: a Multidimensional Psychophysical Study. J. Autism Dev. Disord. 38, 127–137 (2008).

67. L. L. Orefice, A. L. Zimmerman, A. M. Chirila, S. J. Sleboda, J. P. Head, D. D. Ginty, Peripheral Mechanosensory Neuron Dysfunction Underlies Tactile and Behavioral Deficits in Mouse Models of ASDs. Cell 166, 299–313 (2016).

68. H. Oh, S. Lee, Y. Oh, S. Kim, Y. S. Kim, Y. Yang, W. Choi, Y.-E. Yoo, H. Cho, S. Lee, E. Yang, W. Koh, W. Won, R. Kim, C. J. Lee, H. Kim, H. Kang, J. Y. Kim, T. Ku, S.-B. Paik, E. Kim, Kv7/KCNQ potassium channels in cortical hyperexcitability and juvenile seizure-related death in Ank2-mutant mice. Nat. Commun. 14, 3547 (2023).

69. Q. You, Z. Luo, Z. Luo, Y. Kong, Z. Li, J. Yang, X. Li, T. Gao, Involvement of the thalamic reticular nucleus in prepulse inhibition of acoustic startle. Transl. Psychiatry 11, 241 (2021).

70. R. Ayub, K. L. Sun, R. E. Flores, V. T. Lam, B. Jo, M. Saggar, L. K. Fung, Thalamocortical connectivity is associated with autism symptoms in high-functioning adults with autism and typically developing adults. Transl. Psychiatry 11, 93 (2021).

71. G. Avanzini, M. de Curtis, F. Panzica, R. Spreafico, Intrinsic properties of nucleus reticularis thalami neurones of the rat studied in vitro. J. Physiol. 416, 111–122 (1989).

72. L. Cueni, M. Canepari, R. Luján, Y. Emmenegger, M. Watanabe, C. T. Bond, P. Franken, J. P. Adelman, A. Lüthi, T-type Ca2+ channels, SK2 channels and SERCAs gate sleep-related oscillations in thalamic dendrites. Nat. Neurosci. 11, 683–692 (2008).

73. J. R. Huguenard, D. A. McCormick, Thalamic synchrony and dynamic regulation of global forebrain oscillations. Trends Neurosci. 30, 350–356 (2007).

74. M. Steriade, D. A. McCormick, T. J. Sejnowski, Thalamocortical oscillations in the sleeping and aroused brain. Science 262, 679–685 (1993).

75. F. Wang, H. Sun, M. Chen, B. Feng, Y. Lu, M. Lyu, D. Cui, Y. Zhai, Y. Zhang, Y. Zhu, C. Wang, H. Wu, X. Ma, F. Zhu, Q. Wang, Y. Li, The thalamic reticular nucleus orchestrates social memory. Neuron 112, 2368–2385.e11 (2024).

76. C. R. Houser, J. E. Vaughn, R. P. Barber, E. Roberts, GABA neurons are the major cell type of the nucleus reticularis thalami. Brain Res. 200, 341–354 (1980).

77. B. Zikopoulos, H. Barbas, Altered neural connectivity in excitatory and inhibitory cortical circuits in autism. Front. Hum. Neurosci. 7 (2013).

78. F. Filice, L. Janickova, T. Henzi, A. Bilella, B. Schwaller, The Parvalbumin Hypothesis of Autism Spectrum Disorder. Front. Cell. Neurosci. 14, 577525 (2020).

79. V. Marlinski, I. N. Beloozerova, Burst firing of neurons in the thalamic reticular nucleus during locomotion. J. Neurophysiol. 112, 181–192 (2014).

80. J. R. Huguenard, Low-voltage-activated (T-type) calcium-channel genes identified. Trends Neurosci. 21, 451–452 (1998).

81. A. T.-H. Lu, X. Dai, J. A. Martinez-Agosto, R. M. Cantor, Support for calcium channel gene defects in autism spectrum disorders. Mol. Autism 3, 18 (2012).

82. E. M. Talley, L. L. Cribbs, J. H. Lee, A. Daud, E. Perez-Reyes, D. A. Bayliss, Differential distribution of three members of a gene family encoding low voltage-activated (T-type) calcium channels. J. Neurosci. Off. J. Soc. Neurosci. 19, 1895–1911 (1999).

83. S. E. Lee, J. Lee, C. Latchoumane, B. Lee, S.-J. Oh, Z. A. Saud, C. Park, N. Sun, E. Cheong, C.-C. Chen, E.-J. Choi, C. J. Lee, H.-S. Shin, Rebound burst firing in the reticular thalamus is not essential for pharmacological absence seizures in mice. Proc. Natl. Acad. Sci. 111, 11828–11833 (2014).

84. E. K. Harding, A. Dedek, R. P. Bonin, M. W. Salter, T. P. Snutch, M. E. Hildebrand, The T-type calcium channel antagonist, Z944, reduces spinal excitability and pain hypersensitivity. Br. J. Pharmacol. 178, 3517–3532 (2021).

85. A. J. Roebuck, W. N. Marks, M. C. Liu, N. B. Tahir, N. K. Zabder, T. P. Snutch, J. G. Howland, Effects of the T-type calcium channel antagonist Z944 on paired associates learning and locomotor activity in rats treated with the NMDA receptor antagonist MK-801. Psychopharmacology (Berl.) 235, 3339–3350 (2018).

86. W. N. Marks, N. K. Zabder, S. M. Cain, T. P. Snutch, J. G. Howland, The T-type calcium channel antagonist, Z944, alters social behavior in Genetic Absence Epilepsy Rats from Strasbourg. Behav. Brain Res. 361, 54–64 (2019).

87. R. Jagirdar, C.-H. Fu, J. Park, B. F. Corbett, F. M. Seibt, M. Beierlein, J. Chin, Restoring activity in the thalamic reticular nucleus improves sleep architecture and reduces Aβ accumulation in mice. Sci. Transl. Med. 13, eabh4284 (2021).

